# Structure, function and antigenicity of the SARS-CoV-2 spike glycoprotein

**DOI:** 10.1101/2020.02.19.956581

**Authors:** Alexandra C. Walls, Young-Jun Park, M. Alexandra Tortorici, Abigail Wall, Andrew T. McGuire, David Veesler

**Author notes:** These authors contributed equally (Alexandra C. Walls, Young-Jun Park).

## Abstract

The recent emergence of a novel coronavirus associated with an ongoing outbreak of pneumonia (Covid-2019) resulted in infections of more than 72,000 people and claimed over 1,800 lives. Coronavirus spike (S) glycoprotein trimers promote entry into cells and are the main target of the humoral immune response. We show here that SARS-CoV-2 S mediates entry in VeroE6 cells and in BHK cells transiently transfected with human ACE2, establishing ACE2 as a functional receptor for this novel coronavirus. We further demonstrate that the receptor-binding domains of SARS-CoV-2 S and SARS-CoV S bind with similar affinities to human ACE2, which correlates with the efficient spread of SARS-CoV-2 among humans. We found that the SARS-CoV-2 S glycoprotein harbors a furin cleavage site at the boundary between the S_1_/S_2_ subunits, which is processed during biogenesis and sets this virus apart from SARS-CoV and other SARS-related CoVs. We determined a cryo-electron microscopy structure of the SARS-CoV-2 S ectodomain trimer, demonstrating spontaneous opening of the receptor-binding domain, and providing a blueprint for the design of vaccines and inhibitors of viral entry. Finally, we demonstrate that SARS-CoV S murine polyclonal sera potently inhibited SARS-CoV-2 S-mediated entry into target cells, thereby indicating that cross-neutralizing antibodies targeting conserved S epitopes can be elicited upon vaccination.

## INTRODUCTION

Three coronaviruses have crossed the species barrier to cause deadly pneumonia in humans since the beginning of the 21^st^ century: severe acute respiratory syndrome coronavirus (SARS-CoV) (Drosten et al., 2003; Ksiazek et al., 2003), Middle-East respiratory syndrome coronavirus (Zaki et al., 2012) (MERS-CoV) and SARS-CoV-2 (Huang et al., 2020; Zhu et al., 2020). SARS-CoV emerged in the Guangdong province of China in 2002 and spread to five continents through air travel routes, infecting 8,098 people and causing 774 deaths. In 2012, MERS-CoV emerged in the Arabian Peninsula, where it remains a major public health concern, and was exported to 27 countries, infecting a total of ~2,494 individuals and claiming 858 lives. A previously unknown coronavirus, named SARS-CoV-2, was discovered in December 2019 in Wuhan, Hubei Province of China and was sequenced and isolated by January 2020 (Zhou et al., 2020; Zhu et al., 2020). SARS-CoV-2 is associated with an ongoing outbreak of atypical pneumonia (Covid-2019) that has affected over 72,436 people and killed more than 1,868 of them in 29 countries as of February 15^th^ 2020. On January 30^th^ 2020, the World Health Organization declared the SARS-CoV-2 epidemic as a public health emergency of international concern.

MERS-CoV was suggested to originate from bats but the reservoir host fueling spillover to humans is unequivocally dromedary camels (Haagmans et al., 2014; Memish et al., 2013). Both SARS-CoV and SARS-CoV-2 are closely related to each other and originated in bats which most likely serve as reservoir host for these two viruses (Ge et al., 2013; Hu et al., 2017; Li et al., 2005b; Yang et al., 2015a; Zhou et al., 2020). Whereas palm civets and racoon dogs have been recognized as an intermediate host for zoonotic transmission of SARS-CoV between bats and humans (Guan et al., 2003; Kan et al., 2005; Wang et al., 2005), the intermediate host of SARS-CoV-2 remains unknown. The recurrent spillovers of coronaviruses in humans along with detection of numerous coronaviruses in bats, including many SARS-related coronaviruses (SARSr-CoVs), suggest that future zoonotic transmission events may continue to occur (Anthony et al., 2017; Ge et al., 2013; Hu et al., 2017; Li et al., 2005b; Menachery et al., 2015; Menachery et al., 2016; Yang et al., 2015a; Zhou et al., 2020). In addition to the highly pathogenic zoonotic pathogens SARS-CoV, MERS-CoV and SARS-CoV-2, all belonging to the β-coronavirus genus, four low pathogenicity coronaviruses are endemic in humans: HCoV-OC43, HCoV-HKU1, HCoV-NL63 and HCoV-229E.To date, no therapeutics or vaccines are approved against any human-infecting coronaviruses.

Coronavirus entry into host cells is mediated by the transmembrane spike (S) glycoprotein that forms homotrimers protruding from the viral surface (Tortorici and Veesler, 2019). S comprises two functional subunits responsible for binding to the host cell receptor (S_1_ subunit) and fusion of the viral and cellular membranes (S_2_ subunit). For many CoVs, S is cleaved at the boundary between the S_1_ and S_2_ subunits which remain non-covalently bound in the prefusion conformation (Belouzard et al., 2009; Bosch et al., 2003; Burkard et al., 2014; Kirchdoerfer et al., 2016; Millet and Whittaker, 2014, 2015; Park et al., 2016; Walls et al., 2016a). The distal S_1_ subunit comprises the receptor-binding domain(s), and contributes to stabilization of the prefusion state of the membrane-anchored S_2_ subunit that contains the fusion machinery (Gui et al., 2017; Kirchdoerfer et al., 2016; Pallesen et al., 2017; Song et al., 2018; Walls et al., 2016a; Walls et al., 2017b; Yuan et al., 2017). For all CoVs, S is further cleaved by host proteases at the so-called S_2_’ site located immediately upstream of the fusion peptide (Madu et al., 2009; Millet and Whittaker, 2015). This cleavage has been proposed to activate the protein for membrane fusion *via* extensive irreversible conformational changes (Belouzard et al., 2009; Heald-Sargent and Gallagher, 2012; Millet and Whittaker, 2014, 2015; Park et al., 2016; Walls et al., 2017b). As a result, coronavirus entry into susceptible cells is a complex process that requires the concerted action of receptor-binding and proteolytic processing of the S protein to promote virus-cell fusion.

Different coronaviruses use distinct domains within the S_1_ subunit to recognize a variety of attachment and entry receptors, depending on the viral species. Endemic human coronaviruses OC43 and HKU1 attach *via* their S domain A (S^A^) to 5-N-acetyl-9-*O*-acetyl-sialosides found on glycoproteins and glycolipids at the host cell surface to enable entry into susceptible cells (Hulswit et al., 2019; Tortorici et al., 2019; Vlasak et al., 1988). MERS-CoV S, however, uses domain A to engage non-acetylated sialosides as attachment receptors (Li et al., 2017; Park et al., 2019) and promote subsequent binding of domain B (S^B^) to the entry receptor, dipeptidyl-peptidase 4 (Lu et al., 2013; Raj et al., 2013). SARS-CoV and several SARS-related coronaviruses (SARSr-CoV) interact directly with angiotensin-converting enzyme 2 (ACE2) *via* S^B^ to enter target cells (Ge et al., 2013; Kirchdoerfer et al., 2018; Li et al., 2005a; Li et al., 2003; Song et al., 2018; Yang et al., 2015a).

As the coronavirus S glycoprotein is surface-exposed and mediates entry into host cells, it is the main target of neutralizing antibodies (Abs) upon infection and the focus of therapeutic and vaccine design. S trimers are extensively decorated with N-linked glycans that are important for proper folding (Rossen et al., 1998) and to modulate accessibility to host proteases and neutralizing antibodies (Walls et al., 2016b; Walls et al., 2019; Xiong et al., 2017; Yang et al., 2015b). We previously characterized potent human neutralizing Abs from rare memory B cells of individuals infected with SARS-CoV (Traggiai et al., 2004) or MERS-CoV (Corti et al., 2015) in complex with SARS-CoV S and MERS-CoV S to provide molecular-level information of the mechanism of competitive inhibition of S^B^ attachment to the host receptor (Walls et al., 2019). The S230 anti-SARS-CoV antibody also acted by functionally mimicking receptor-attachment and promoting spike fusogenic conformational rearrangements through a ratcheting mechanism that elucidated the unique nature of the coronavirus membrane fusion activation (Walls et al., 2019).

We report here that ACE2 could mediate SARS-CoV-2 S-mediated entry into cells, establishing it as a functional receptor for this newly emerged coronavirus. The SARS-CoV-2 S^B^ engages human ACE2 (hACE2) with comparable affinity than SARS-CoV S^B^ from viral isolates associated with the 2002-2003 epidemic (i.e. binding with high affinity to hACE2). Tight binding to hACE2 could explain the efficient transmission of SARS-CoV-2 in humans, as was the case for SARS-CoV. We identified the presence of an unexpected furin cleavage site at the S_1_/S_2_ boundary of SARS-CoV-2 S, which is cleaved during biosynthesis, a novel feature setting this virus apart from SARS-CoV and SARSr-CoVs. Abrogation of this cleavage motif moderately affected SARS-CoV-2 S-mediated entry into VeroE6 or BHK cells but may contribute to expand the tropism of this virus, as reported for several highly pathogenic avian influenza viruses and pathogenic Newcastle disease virus (Klenk and Garten, 1994; Steinhauer, 1999). We determined a cryo-electron microscopy structure of the SARS-CoV-2 S ectodomain trimer and reveal that it adopts multiple S^B^ conformations that are reminiscent of previous reports on both SARS-CoV S and MERS-CoV S. Finally, we show that SARS-CoV S mouse polyclonal sera potently inhibited entry into target cells of SARS-CoV-2 S pseudotyped viruses. Collectively, these results pave the way for designing vaccines eliciting broad protection against SARS-CoV-2, SARS-CoV and SARSr-CoV.

## RESULTS

### ACE2 is an entry receptor for SARS-CoV-2

The SARS-CoV-2 S glycoprotein shares ~80% amino acid sequence identity with the SARS-CoV S Urbani and with bat SARSr-CoV ZXC21 S and ZC45 S glycoprotein. The latter two SARSr-CoV sequences were identified from *Rinolophus sinicus* (Chinese horseshoe bats), the species from which SARSr-CoV WIV-1 and WIV-16 were isolated (Ge et al., 2013; Yang et al., 2015a). Furthermore, Zhou *et al* recently reported that SARS-CoV-2 is most closely related to the bat SARSr-CoV RaTG13 with which it forms a distinct lineage from other SARSr-CoVs, and that their S glycoproteins share 98% amino acid sequence identity (Zhou et al., 2020). SARS-CoV recognizes its entry receptor human ACE2 (hACE2) at the surface of type II pneumocytes, using S^B^ which shares ~75% overall amino acid sequence identity with SARS-CoV-2 S^B^ and 50% identity within their receptor-binding motifs (RBMs) (Li et al., 2005a; Li et al., 2003; Li et al., 2005c; Wan et al., 2020). Previous studies also showed that the host proteases cathepsin L and TMPRSS2 prime SARS-CoV S for membrane fusion through cleavage at the S_1_/S_2_ and at the S_2_’ sites (Belouzard et al., 2009; Bosch et al., 2008; Glowacka et al., 2011; Matsuyama et al., 2010; Millet and Whittaker, 2015; Shulla et al., 2011).

We set out to investigate the functional determinants of S-mediated entry into target cells using a murine leukemia virus (MLV) pseudotyping system (Millet and Whittaker, 2016). To assess the ability of SARS-CoV-2 S to promote entry into target cells, we first compared transduction of SARS-CoV-2 S-MLV and SARS-CoV S-MLV into VeroE6 cells, that are known to express ACE2 and support SARS-CoV replication (Drosten et al., 2003; Ksiazek et al., 2003). Both pseudoviruses entered cells equally well **(Fig. 1 A)**, suggesting that SARS-CoV-2 S-MLV could potentially use African green monkey ACE2 as entry receptor. To confirm these results, we evaluated entry into BHK cells and observed that transient transfection with hACE2 rendered them susceptible to transduction with SARS-CoV-2 S-MLV **(Fig. 1 B)**. These results demonstrate hACE2 is a functional receptor for SARS-CoV-2, in agreement with recently reported findings (Hoffmann et al., 2020; Letko and Munster, 2020; Zhou et al., 2020).

**Figure 1.**
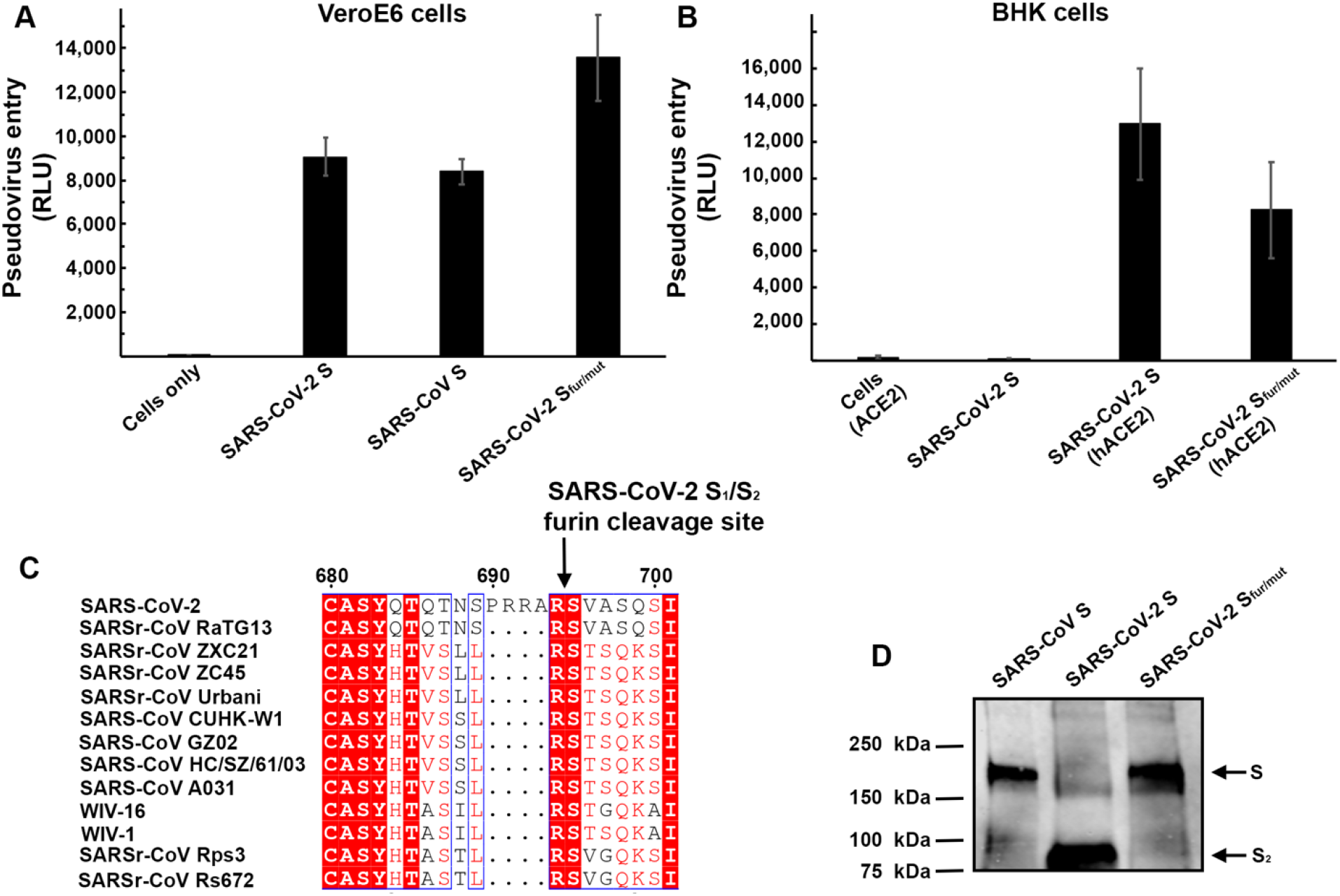
hACE2 is a functional receptor for SARS-CoV-2 S. **A.** Entry of MLV pseudotyped with SARS-CoV-2 S, SARS-CoV-2 S_fur/mut_ and SARS-CoV S in VeroE6 cells. **B.** Entry of MLV pseudotyped with SARS-CoV-2 S or SARS-CoV-2 S_fur/mut_ in BHK cells transiently transfected with hACE2. The experiments were carried out in triplicate with two independent pseudovirus preparations and a representative experiment is shown. **C.** Sequence alignment of SARS-CoV-2 S with multiple related SARS-CoV and SARSr-CoV S glycoproteins reveals the introduction of an S_1_/S_2_ furin cleavage site in this novel coronavirus. Identical and similar positions are respectively shown with white or red font. The four amino acid residue insertion at SARS-CoV-2 S positions 690-693 is indicated with periods. The entire sequence alignment is presented in Fig. S1. **D.** Western blot analysis of SARS-CoV-2 S-MLV, SARS-CoV-2 S_fur/mut_-MLV and SARS-CoV S-MLV pseudovirions using an anti-SARS-CoV S_2_ antibody.

Sequence analysis of SARS-CoV-2 S reveals the presence of a four amino acid residue insertion at the boundary between the S_1_ and S_2_ subunits compared to SARS-CoV S and SARSr-CoV S **(Fig. 1 C)**. This results in the introduction of a furin cleavage site, a feature conserved among the 103 SARS-CoV-2 isolates sequenced to date but not in the closely related RaTG13 S (Zhou et al., 2020). Using Western blot analysis, we observed that SARS-CoV-2 S was virtually entirely processed at the S_1_/S_2_ site during biosynthesis in HEK293T cells, presumably by furin in the Golgi compartment **(Fig. 1 D)**. This observation contrasts with SARS-CoV S which was incorporated into pseudovirions largely uncleaved **(Fig. 1 D)**. To study the influence on pseudovirus entry of the SARS-CoV-2 S_1_/S_2_ furin cleavage site, we designed an S mutant lacking the four amino acid residue insertion and the furin cleavage site by mutating Q_686_T**NSPRR**AR↓SV_696_ (wildtype SARS-CoV-2 S) to Q_686_T**IL**R↓SV_692_ (SARS-CoV-2 S_fur/mut_). SARS-CoV-2 S_fur/mut_ preserves only the conserved Arg residue at position 994 of wildtype SARS-CoV-2 S thereby mimicking the S_1_/S_2_ cleavage site of the related SARSr-CoV S CZX21 **(Fig. 1 D).** SARS-CoV-2 S_fur/mut_ is therefore expected to undergo processing at the S_1_/S_2_ site upon encounter of a target cell, similarly to SARS-CoV S and SARSr-CoV S (i.e. via TMPRSS2 and/or cathepsin L). As expected, SARS-CoV-2 S_fur/mut_-MLV harbored uncleaved S upon budding **(Fig. 1 D)**. The observed transduction efficiency of VeroE6 cells was higher for SARS-CoV-2 S_fur/mut_-MLV than for SARS-CoV-2 S-MLV **Fig. 1 A)** whereas the opposite trend was observed for transduction of hACE2-expressing BHK cells **(Fig. 1 B)**. These results suggest that S_1_/S_2_ cleavage during S biosynthesis was not necessary for S-mediated entry in the conditions of our experiments **(Fig. 1 C-D)**. We speculate that the detection of a polybasic cleavage site in the fusion glycoprotein of SARS-CoV-2 could putatively expand its tropism and/or enhance its transmissibility, compared to SARS-CoV and SARSr-CoV isolates, due to the near-ubiquitous distribution of furin-like proteases and their reported effects on other viruses (Klenk and Garten, 1994; Millet and Whittaker, 2015; Steinhauer, 1999).

### SARS-CoV-2 recognizes human ACE2 with comparable affinity than SARS-CoV

The binding affinity of SARS-CoV for hACE2 correlate with the overall rate of viral replication in distinct species, transmissibility and disease severity (Guan et al., 2003; Li et al., 2004; Li et al., 2005c; Wan et al., 2020). Indeed, specific S^B^ mutations enabled efficient binding to hACE2 of SARS-CoV isolates from the three phases of the 2002-2003 epidemic, which were associated with marked disease severity (Consortium, 2004; Kan et al., 2005; Li et al., 2005c; Sui et al., 2004). In contrast, SARS-CoV isolates detected during the brief 2003-2004 re-emergence interacted more weakly with hACE2, but tightly with civet ACE2, and had low pathogenicity and transmissibility (Consortium, 2004; Kan et al., 2005; Li et al., 2005c).

To understand the contribution of receptor interaction to the infectivity of SARS-CoV-2, we characterized engagement of hACE2 by SARS-CoV-2 S^B^ and SARS-CoV S^B^ side-by-side. We used biolayer interferometry to study binding kinetics and affinity of the purified hACE2 ectodomain to SARS-CoV-2 S^B^ and SARS-CoV S^B^ immobilized at the surface of biosensors. We found that hACE2 bound to SARS-CoV-2 S^B^ and SARS-CoV S^B^ with respective equilibrium dissociation constants of 2.9 nM **(Fig. 2 A)** and 7.7 nM **(Fig. 2 B)**, and comparable kinetic rate constants although the off-rate was slightly higher for SARS-CoV S^B^ **(Table 1)**. Previous structural work identified 14 positions that are key for binding of SARS-CoV S^B^ to hACE2: T402, R426, Y436, Y440, Y442, L472, N473, Y475, N479, Y484, T486, T487, G488 and Y491 (Li et al., 2005a). Analysis of the 104 SARS-CoV-2 genome sequences available from GISAID (Elbe and Buckland-Merrett, 2017) shows that 8 out of these 14 positions are strictly conserved in SARS-CoV-2 S^B^ whereas the other 6 positions are (semi)-conservatively substituted: R426_SARS-CoV_N448_SARS-CoV-2_, Y442_SARS-CoV_L464_SARS-CoV-2_, L472_SARS-CoV_F495_SARS-CoV-2_, N479_SARS-CoV_Q502_SARS-CoV-2_, Y484_SARS-CoV_Q507_SARS-CoV-2_, T487_SARS-CoV_N510_SARS-CoV-2_ **(Fig. 2 C)**. The conservation of many key contact residues could explain the similar binding affinities of SARS-CoV-2 S^B^ and SARS-CoV S^B^ for hACE2. No mutations of residues predicted to contact hACE2 have been observed among SARS-CoV-2 S sequences available to date. However, we note that SARSr-CoV ZXC21 and ZC45, which harbor the most closely related S sequences after SARSr-CoV RaTG13, comprise two deletions in the RBD that could affect binding to hACE2 (between SARS-CoV-2 S residues 482-489 and 494-499) **(Fig. S 1)**. Collectively, these results suggest that SARS-CoV-2 is at least as well adapted to the hACE2 orthologue as the 2002-2003 epidemic strains of SARS-CoV, which could explain the efficient transduction efficiency mediated by their respective S glycoproteins **(Fig. 1 A-B)** and the current rapid SARS-CoV-2 transmission in humans.

**Figure 2.**
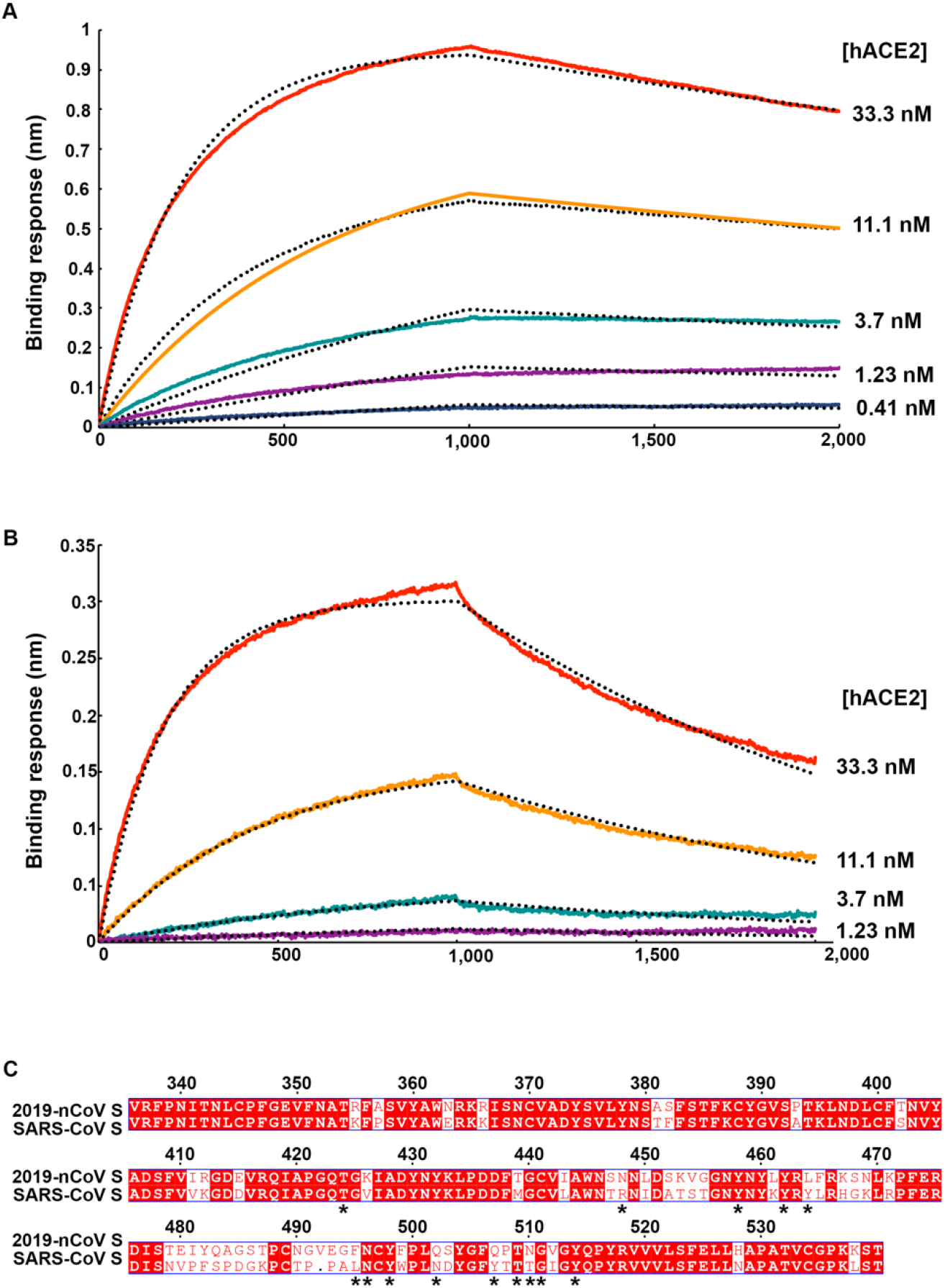
SARS-CoV-2 S recognizes hACE2 with comparable affinity to SARS-CoV S. **A-B.** Biolayer interferometry binding analysis of the hACE2 ectodomain to immobilized SARS-CoV-2 S^B^ (B) or SARS-CoV S^B^ (C). The experiments were performed in duplicates with different protein preparations and one representative set of curves is shown. Dotted lines correspond to a global fit of the data using a 1:1 binding model. **C.** Sequence alignment of SARS-CoV-2 S^B^ and SARS-CoV S^B^ Urbani (late phase of the 2002-2003 SARS-CoV epidemic). Identical and similar positions are respectively shown with white or red font. The single amino acid insertion at position 492 of the SARS-CoV-2 S^B^ is indicated with a period at the corresponding SARS-CoV S^B^ position. The 14 residues that are key for binding of SARS-CoV S^B^ to hACE2 are labeled with a star.

**Table 1.** Kinetic analysis of hACE2 binding to SARS-CoV-2 S^B^ and SARS-CoV S^B^ by biolayer interferometry. Values reported represent the global fit to the data shown in Fig. 2 A-B and the averages obtained from 5 (SARS-CoV-2) or 4 (SARS-CoV) replicates carried out with different protein preparations.

|  | **SARS-CoV-2 S<sup>B</sup>** | **SARS-CoV-2 S<sup>B</sup>**<br>**Average** | **SARS-CoV S<sup>B</sup>** | **SARS-CoV S<sup>B</sup>**<br>**Average** |
| --- | --- | --- | --- | --- |
| **K<sub>D</sub> (nM)** | 1.2 +/-<br>0.1 | 2.9 +/- 3.6 | 4.9 +/- 0.1 | 7.7 +/- 8.8 |
| **k<sub>on</sub>** | 1.4 x 10<sup>5</sup> | 2.3 +/- 1.4 x 10<sup>5</sup> | 1.4 x 10<sup>5</sup> | 1.7 +/- 7.1 x 10<sup>5</sup> |
| **(M<sup>-1</sup>.s<sup>-1</sup>)** |  |  |  |  |
| **k<sub>off</sub> (s<sup>-1</sup>)** | 1.6 x 10<sup>-4</sup> | 1.7 +/- 0.84 x 10<sup>-4</sup> | 7.61x 10<sup>-4</sup> | 8.7 +/- 5.1 x 10<sup>-4</sup> |

### Architecture of the SARS-CoV-2 spike glycoprotein trimer

To enable single-particle cryoEM study of the SARS-CoV-2 S glycoprotein, we designed a prefusion stabilized ectodomain trimer construct with an abrogated furin S_1_/S_2_ cleavage site (Tortorici et al., 2019; Walls et al., 2017a; Walls et al., 2016a; Walls et al., 2019), two consecutive proline stabilizing mutations (Kirchdoerfer et al., 2018; Pallesen et al., 2017) and a C-terminal foldon trimerization domain (Miroshnikov et al., 1998). 3D classification of the cryoEM data revealed the presence of multiple conformational states of SARS-CoV-2 S corresponding to distinct organization of the S^B^ domains within the S_1_ apex. Approximately half of the particle images selected correspond to trimers harboring a single S^B^ domain opened whereas the remaining half was accounted for by closed trimers with the 3 S^B^ domains closed. The observed conformational variability of S^B^ domains is reminiscent of observations made with SARS-CoV S and MERS-CoV S trimers although we did not detect trimers with two S^B^ domains open and the distribution of particles across the S conformational landscape varies among studies (Gui et al., 2017; Kirchdoerfer et al., 2018; Pallesen et al., 2017; Song et al., 2018; Walls et al., 2019; Yuan et al., 2017).

We determined a reconstruction of the closed SARS-CoV-2 S ectodomain trimer at 3.3 Å resolution (applying 3-fold symmetry) and an asymmetric reconstruction of the trimer with a single S^B^ domain opened at 3.7 Å resolution **(Fig 3 A-H, Fig S2, Table 2)**. The S_2_ fusion machinery is the best resolved part of the map whereas the S^A^ and S^B^ domains are less well resolved, presumably due to conformational heterogeneity. The final atomic model comprises residues 36-1157, with internal breaks corresponding to flexible regions, and lacks the C-terminal most segment (including the heptad repeat 2) which is not visible in the map, as is the case for all S structures determined to date.

**Figure 3.**
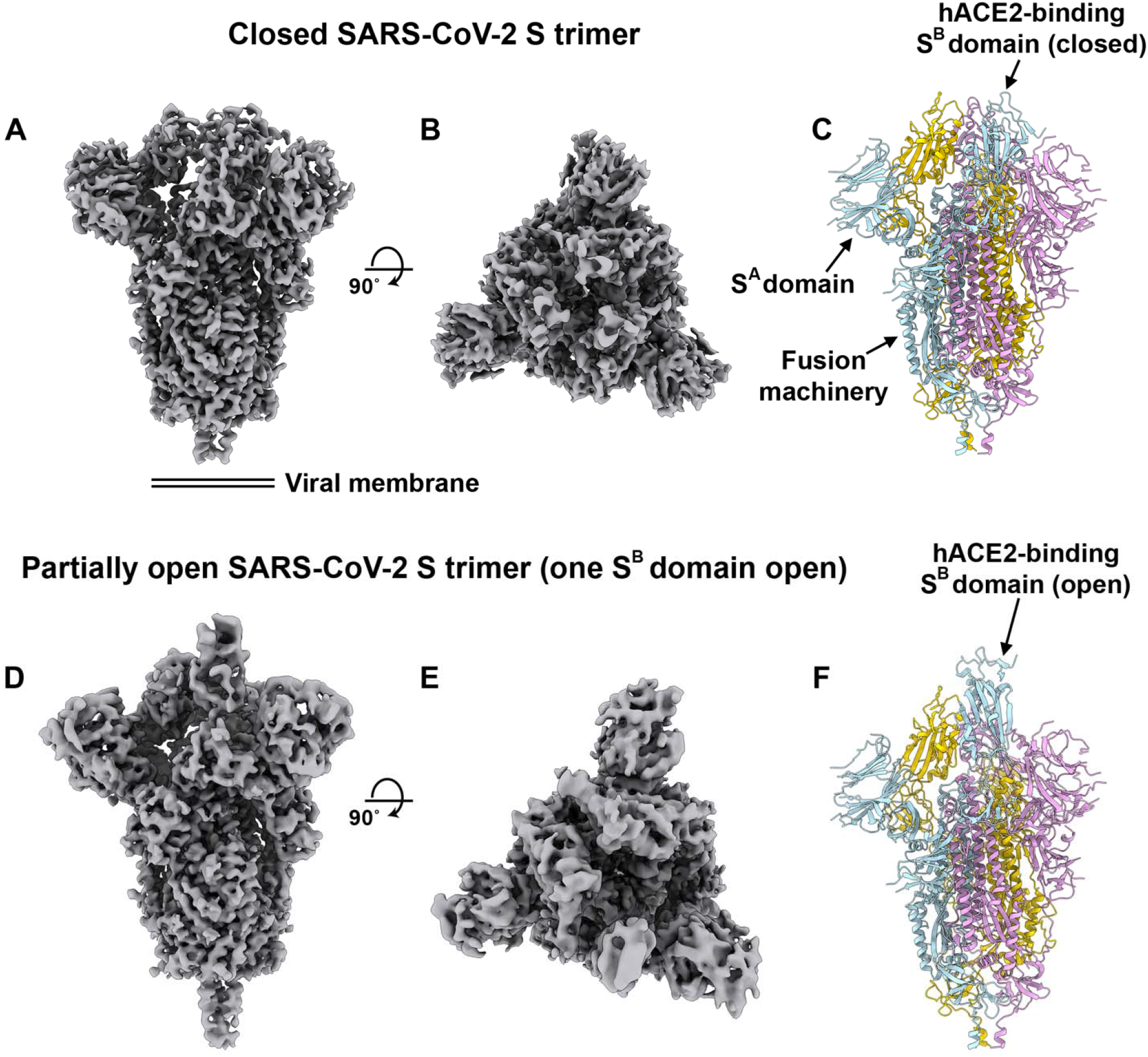
CryoEM structures of the SARS-CoV-2 S glycoprotein. **A-B.** Two orthogonal views of the closed SARS-CoV-2 S trimer cryoEM map. **C.** Atomic model of the closed SARS-CoV-2 S trimer in the same orientation as in panel A. **D-E.** Two orthogonal views of the partially open SARS-CoV-2 S trimer cryoEM map (one S^B^ domain is open). **F.** Atomic model of the closed SARS-CoV-2 S trimer in the same orientation as in panel D. The glycans were omitted for clarity.

**Table 2.** CryoEM data collection and refinement statistics

|  | SARS-CoV-2<br>S (one S<sup>B</sup><br>open) | SARS-CoV-2 S<br>(pseudo-<br>closed) |
| --- | --- | --- |
| **Data collection and processing** |  |  |
| Magnification | 130,000 | 130,000 |
| Voltage (kV) | 300 | 300 |
| Electron exposure (e<sup>-</sup>/Å<sup>2</sup>) | 70 | 70 |
| Defocus range (μm) | 0.5-3.0 | 0.5-3.0 |
| Pixel size (Å) | 0.525 | 0.525 |
| Symmetry imposed | C1 | C1 |
| Final particle images (no.) | 115,591 | 99,530 |
| Map resolution (Å) | 3.3 | 3.8 |
| FSC threshold | 0.143 | 0.143 |
| Map sharpening *B* factor (Å<sup>2</sup>) | -136.8 | -136.8 |
| **Validation** |  |  |
| MolProbity score | 1.23 |  |
| Clashscore | 1.66 |  |
| Poor rotamers (%) | 0.24 |  |
| Ramachandran plot |  |  |
| Favored (%) | 97.57 |  |
| Allowed (%) | 99.76 |  |
| Disallowed (%) | 0.24 |  |
| EMRinger Score | 2.75 |  |

Overall, the SARS-CoV-2 S ectodomain is a 160Å-long trimer with a triangular crosssection, resembling the closely related SARS-CoV S structure **(Fig S3 A-F)**(Gui et al., 2017; Kirchdoerfer et al., 2018; Song et al., 2018; Walls et al., 2019; Yuan et al., 2017).

As is the case for other β-coronavirus S glycoproteins, including SARS-CoV S (Gui et al., 2017; Kirchdoerfer et al., 2018; Pallesen et al., 2017; Park et al., 2019; Song et al., 2018; Tortorici et al., 2019; Walls et al., 2016a; Walls et al., 2016b; Walls et al., 2019; Yuan et al., 2017), the S_1_ subunit has a V-shaped architecture **(Fig S3 B-E)**. In the closed S trimer, the three hACE2-recognition motifs, whose location was inferred based on the crystal structure of SARS-CoV S^B^ in complex with hACE2 (Li et al., 2005a), are buried at the interface between protomers. As a result, SARS-CoV-2 S^B^ opening is expected to be necessary for interacting with ACE2 at the host cell surface and initiating the conformational changes leading to cleavage of the S_2_’ site, membrane fusion and viral entry (Gui et al., 2017; Kirchdoerfer et al., 2018; Millet and Whittaker, 2014; Pallesen et al., 2017; Park et al., 2016; Song et al., 2018; Walls et al., 2019; Yuan et al., 2017).

As the SARS-CoV-2 and SARS-CoV S_2_ subunits share 88% sequence identity, they are structurally conserved and can be superimposed with 1.2 Å rmsd over 417 aligned Cα positions **(Fig S3 E-F)**. Only the most N-terminal part of the fusion peptide is resolved in the map, as was the case for previously determined SARS-CoV S structures. The sequence and conformational conservation of the fusion peptide region observed across SARS-CoV-2 S and SARS-CoV S suggests that antibodies targeting this functionally important motif might cross-react and neutralize the two viruses as well as related coronaviruses **(Fig S3 G-H)**

We previously showed that coronavirus S glycoproteins are densely decorated by heterogeneous N-linked glycans protruding from the trimer surface (Walls et al., 2016b; Walls et al., 2019; Xiong et al., 2017). These oligosaccharides participate in S folding (Rossen et al., 1998), they affect priming by host proteases (Yang et al., 2015b) and modulate antibody recognition (Pallesen et al., 2017; Walls et al., 2019). SARS-CoV-2 S comprise 22 N-linked glycosylation sequons per protomer and 16 of them are resolved in the cryoEM map **(Fig. 4)**. By comparison, SARS-CoV S possesses 23 N-linked glycosylation sequons per protomer and we previously confirmed experimentally that at least 19 of them are glycosylated (Walls et al., 2019). 20 out of 22 SARS-CoV-2 S N-linked glycosylation sequons are conserved in SARS-CoV S **(Table 3)**. Specifically, 9 out of 13 glycans in the S_1_ subunit and all 9 glycans in the S_2_ subunit are conserved among SARS-CoV-2 S and SARS-CoV S. Furthermore, S_2_ oligosaccharides are mostly conserved across SARSr-CoV S glycoproteins, suggesting that accessibility of the fusion machinery to antibodies will be comparable among these viruses.

**Figure 4.**
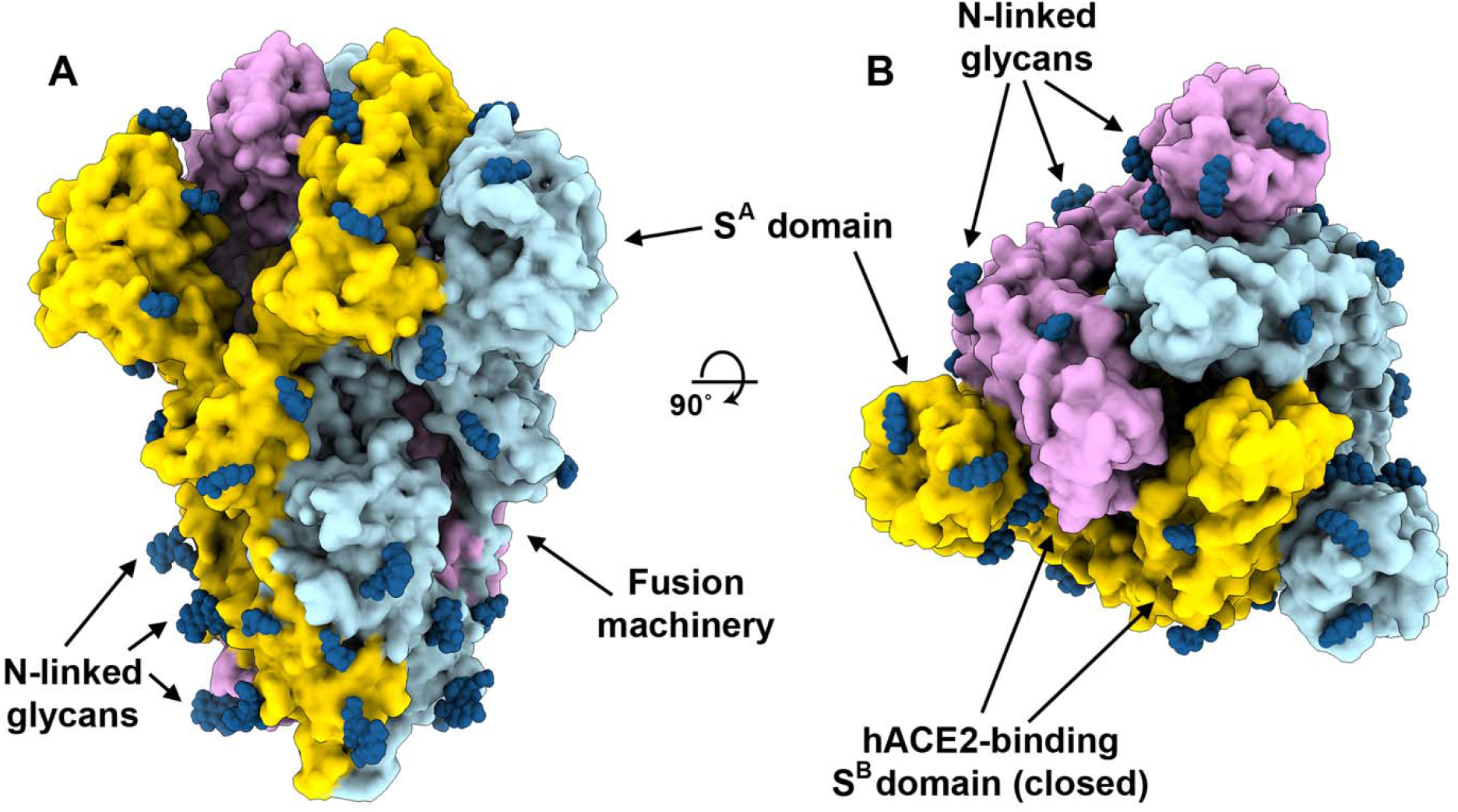
Organization of the SARS-CoV-2 S glycan shield. **A-B.** Two orthogonal views of the SARS-CoV-2 S structure rendered as a surface with glycans rendered as dark blue spheres.

**Table 3.** Conservation of N-linked glycosylation sequons in SARS-CoV-2 S and SARS-CoV S. Red indicates the absence of a glycosylation sequon and deletions are indicated with periods. Glycans observed in the SARS-CoV-2 S cryoEM map are underlined.

| SARS-CoV-2 S | SARS-CoV S |
| --- | --- |
| N<sub>26</sub>LT | T<sub>21</sub>FD |
| P<sub>34</sub>PA | N<sub>29</sub>YT |
| <u>N<sub>70</sub>VT</u> | N<sub>65</sub>VT |
| H<sub>78</sub>VS | N<sub>73</sub>HT |
| N<sub>83</sub>GT | ... |
| S<sub>121</sub>KT | N<sub>109</sub>KS |
| N<sub>130</sub>NA | N<sub>118</sub>NS |
| <u>N<sub>131</sub>AT</u> | N<sub>119</sub>ST |
| N<sub>158</sub>KS | T<sub>146</sub>QT |
| <u>N<sub>174</sub>CT</u> | N<sub>158</sub>CT |
| <u>N<sub>243</sub>IT</u> | N<sub>227</sub>IT |
| <u>N<sub>291</sub>GT</u> | N<sub>269</sub>GT |
| <u>N<sub>340</sub>IT</u> | N<sub>318</sub>IT |
| <u>N<sub>352</sub>AT</u> | N<sub>330</sub>AT |
| N<sub>379</sub>SA | N<sub>357</sub>ST |
| <u>N<sub>612</sub>TS</u> | N<sub>589</sub>AS |
| <u>N<sub>625</sub>CT</u> | N<sub>602</sub>CT |
| <u>N<sub>666</sub>NS</u> | D<sub>643</sub>TS |
| <u>N<sub>718</sub>NS</u> | N<sub>691</sub>NT |
| <u>N<sub>726</sub>FT</u> | N<sub>699</sub>FS |
| <u>N<sub>810</sub>FS</u> | N<sub>783</sub>FS |
| <u>N<sub>1083</sub>FT</u> | N<sub>1056</sub>FT |
| <u>N<sub>1107</sub>GT</u> | N<sub>1080</sub>GT |
| <u>N<sub>1143</sub>NT</u> | N<sub>1116</sub>NT |
| N<sub>1167</sub>HT | N<sub>1140</sub>HT |
| N<sub>1182</sub>AS | N<sub>1155</sub>AS |
| N<sub>1203</sub>ES | N<sub>1176</sub>ES |

### SARS-CoV S elicits neutralizing antibodies against SARS-CoV-2 S

Mapping of the sequence conservation across multiple sarbecovirus S sequences underscores that the S_2_ fusion machinery is more conserved than the S_1_ subunit with the highest divergence found within S^A^ and S^B^ **(Fig. 5A-B)**. These observations are in line with (i) the fact that some, but not all of these viruses use ACE2 as entry receptor (Ge et al., 2013; Ren et al., 2008; Yang et al., 2015a); and (ii) that the S_1_ subunit is more exposed at the viral surface than the fusion machinery, and is likely to be subject to a more stringent selection pressure from the immune system.

**Figure 5.**
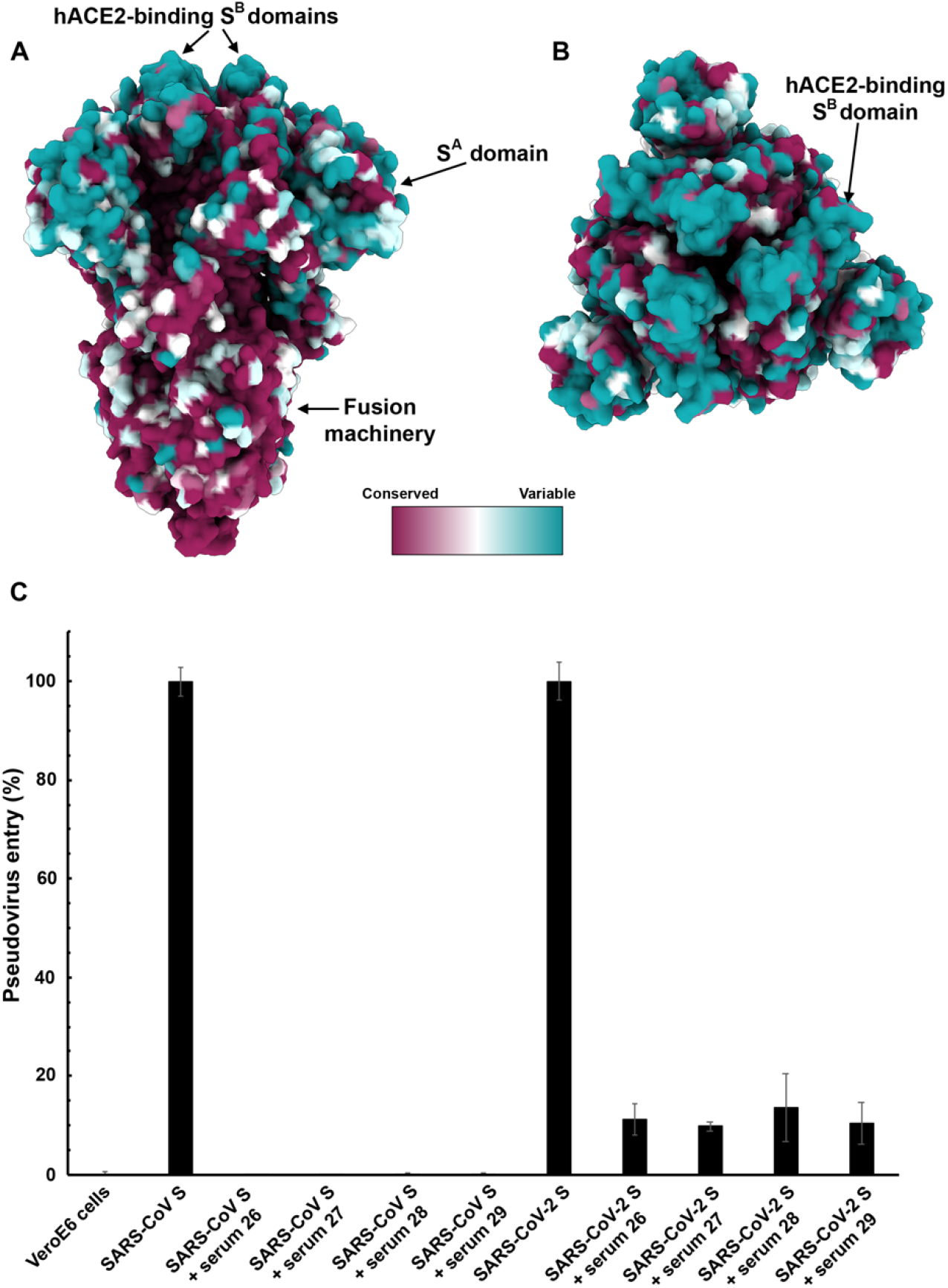
SARS-CoV S elicits antibodies neutralizing SARS-CoV-2 S mediated entry into host cells. **A-B.** Sequence conservation of sarbecovirus S glycoproteins plotted on the SARS-CoV-2 S structure. **C.** Entry of SARS-CoV-2 and SARS-CoV S pseudotyped viruses is potently inhibited by four SARS-CoV S mouse polyclonal immune plasma.

Based on these observations, we hypothesized that exposure to one of the two viruses could elicit cross-reactive and potentially neutralizing antibodies against the other virus. We therefore investigated the ability of plasma from four mice immunized with a stabilized SARS-CoV S to inhibit SARS-CoV-2 S- and SARS-CoV S-mediated entry into target cells. All sera tested completely inhibited transduction of SARS-CoV S-MLV and reduced SARS-CoV-2 S-MLV transduction to ~10% of control in VeroE6 cells **(Fig. 5C)**. The elicitation of a heterotypic response blocking SARS-CoV-2 S-mediated entry into host cells concurs with the sequence and structural conservation of SARS-CoV-2 S and SARS-CoV S along with their comparable glycans shields, and suggests that immunity against one virus of the sarbecovirus subgenus can potentially provide protection against related viruses.

## DISCUSSION

Receptor recognition is the first step of viral infection and is a key determinant of host cell and tissue tropism. Enhanced binding affinity between SARS-CoV S and hACE2 was proposed to correlate with increased virus transmissibility and disease severity in humans (Li et al., 2005c). Indeed, SARS-CoV isolates from the three phases of the 2002-2003 epidemic were more efficiently transmitted among humans and more pathogenic than the isolates associated with the 2003-2004 re-emergence that caused only a few cases, in line with their binding affinities for hACE2 (Consortium, 2004; Kan et al., 2005; Li et al., 2005c). Moreover, the ability to engage ACE2 from different animal species appears to reflect host susceptibility to SARS-CoV infection and facilitated the jump of the virus from animals to humans (Li, 2008; Li et al., 2004). We report here that SARS-CoV-2 uses hACE2 as entry receptor to which it binds with similar affinity than the 2002-2003 SARS-CoV isolates which suggests it can spread efficiently in humans, in agreement with the numerous SARS-CoV-2 human-to-human transmission events reported to date.

Besides binding to host cell receptors, priming of the S glycoprotein by host proteases through cleavage at the S_1_/S_2_ and the S_2_’ sites is another crucial factor modulating tropism and pathogenicity (Millet and Whittaker, 2015). For instance, entry of the MERS-CoV-related bat coronavirus HKU4 into human cells required addition of exogenous trypsin, indicating that S proteolytic activation of this bat virus did not occur in human cells (despite its ability to recognize human DPP4)(Wang et al., 2014; Yang et al., 2014). Subsequent work suggested that a glycan present near the S_1_/S_2_ boundary accounted for the lack of proteolytic priming of HKU4 S and that its removal enhanced pseudovirus entry in human cells (Yang et al., 2015b). The presence of a polybasic cleavage site, that can be processed by furin-like proteases, is a signature of several highly pathogenic avian influenza viruses and pathogenic Newcastle disease virus (Klenk and Garten, 1994; Steinhauer, 1999). Strikingly, SARS-CoV-2 S harbors a furin cleavage site at the S_1_/S_2_ boundary, which is processed during biosynthesis. The presence of a furin cleavage site sets SARS-CoV-2 S apart from SARS-CoV S (and SARSr-CoV S) that possesses a monobasic S_1_/S_2_ cleavage site processed upon entry of target cells (Belouzard et al., 2009; Bosch et al., 2008; Glowacka et al., 2011; Matsuyama et al., 2010; Millet and Whittaker, 2015; Shulla et al., 2011). We speculate that the almost ubiquitous expression of furin-like proteases could participate in expanding SARS-CoV-2 cell and tissue tropism, relative to SARS-CoV, as well as increase its transmissibility and/or alter its pathogenicity.

We previously suggested coronaviruses use conformational masking and glycan shielding to limit recognition by the immune response of infected hosts (Walls et al., 2016b; Walls et al., 2019; Xiong et al., 2017). Similarly to SARS-CoV S and MERS-CoV S (Gui et al., 2017; Kirchdoerfer et al., 2018; Pallesen et al., 2017; Song et al., 2018; Walls et al., 2019; Yuan et al., 2017), we found that the SARS-CoV-2 S trimer exist in multiple, distinct conformational states resulting from S^B^ opening at the trimer apex. These structural changes are necessary for receptor engagement of these three viruses and lead to initiation of fusogenic conformational changes (Song et al., 2018; Walls et al., 2017b; Walls et al., 2019). In contrast, only closed S trimers have been detected for HCoV-NL63 (Walls et al., 2016b), HCoV-OC43 (Tortorici et al., 2019), HCoV-HKU1 (Kirchdoerfer et al., 2016) and HCoV-229E (Li et al., 2019). As HCoV-NL63 and HCoV-229E are known to engage protein receptors through S^B^ (Wong et al., 2017; Wu et al., 2009), trimer opening is also expected to occur to expose their receptor-recognition motifs that are otherwise buried at the interface between protomers in the closed S trimer (Li et al., 2019; Walls et al., 2016b). Regardless of the nature of the receptor and of the location of the receptor-binding domains, removal of the trimeric S_1_ crown is expected to be necessary for all coronaviruses to allow the large-scale S_2_ conformational changes leading to fusion of the viral and host membranes (Walls et al., 2017b). Collectively, these data underscores that S glycoprotein trimers found in highly pathogenic human coronaviruses appear to exist in partially opened states whereas they remain largely closed in human coronaviruses associated with common colds. Based on the aforementioned data correlating the binding affinity of SARS-CoV for hACE2 with the rate of transmissibility, viral replication in distinct species and disease severity (Guan et al., 2003; Li et al., 2004; Li et al., 2005c; Wan et al., 2020), we hypothesize that the most pathogenic coronaviruses will exhibit S glycoprotein trimers spontaneously sampling closed and open conformations, as is the case for SARS-CoV-2, SARS-CoV and MERS-CoV.

The striking structural similarity and sequence conservation among the SARS-CoV-2 S and SARS-CoV S glycoproteins emphasize the close relationship between these two viruses that recognize hACE2 to enter target cells. This resemblance is further strengthened by our finding that SARS-CoV S elicited polyclonal Ab responses potently neutralizing SARS-CoV-2 S-mediated entry into cells. We surmise most of these Abs target the highly conserved S_2_ subunit (including the fusion peptide region) based on its structural similarity across SARS-CoV-2 and SARS-CoV and previous reports showing that sera from SARS-CoV-infected individuals target this region (Zhang et al., 2004). We note that most SARS-CoV neutralizing Abs isolated to date target the S^B^ domain and several of them recognize the RBM and prevent receptor engagement (Hwang et al., 2006; Rockx et al., 2008; Rockx et al., 2010; Traggiai et al., 2004; Walls et al., 2019). As the SARS-CoV-2 and SARS-CoV S^B^ domains share 75% amino acid sequence identity, future work will be necessary to evaluate how many of these antibodies neutralize the newly emerged coronavirus. These findings also indicate that it might be difficult to distinguish exposure to SARS-CoV-2 from other SARSr-CoVs in serological studies using S ectodomain trimers and that specific assays will need to be designed. Our results provide a structural framework to identify conserved and accessible epitopes across S glycoproteins that will support ongoing vaccine design efforts. Finally, elicitation of diverse, polyclonal Ab responses might prove key in light of the diversity of viruses circulating in animal reservoirs and to prevent the possible emergence of viral neutralization escape mutants.

## ACKNOWLEDGEMENTS

This study was supported by the National Institute of General Medical Sciences (R01GM120553, D.V.), the National Institute of Allergy and Infectious Diseases (HHSN272201700059C, DV), a Pew Biomedical Scholars Award (D.V.), an Investigators in the Pathogenesis of Infectious Disease Award from the Burroughs Wellcome Fund (D.V.), the Open Philantropic Foundation (D.V.) and the Pasteur Institute (M.A.T.). We are grateful to Lynda Stuart for sharing the full-length hACE2 plasmid, to Gary Whittaker for the MLV pseudotyping system and to Ning Zheng for providing access to the Octet device.

## AUTHOR CONTRIBUTIONS

A.C.W., Y.J.P, A.T.M, and D.V. designed the experiments. A.C.W. and M.A.T. expressed and purified the proteins. A.C.W. carried out binding assays and pseudovirus entry assays. Y.J.P. prepared samples for cryoEM and collected the data. Y.J.P. and D.V. processed the data, built and refined the atomic models. A.W. and A.T.M. immunized mice. A.C.W., Y.J.P and D.V. analyzed the data and prepared the manuscript with input from all authors.

## MATERIALS AND METHODS

### Transient expression of SARS-CoV-2 and SARS-CoV S^B^

The SARS-CoV S^B^ construct was cloned from a SARS-CoV S ectodomain (Walls et al., 2019) synthesized by GeneArt (ThermoFisher Scientific) into a modified pOPING vector with an N-terminal mu-phosphatase signal peptide and a C-terminal hexa-histidine tag (G-HHHHHH). The boundaries of the construct are N-terminal _306_RVVPSG_311_ and C-terminal _571_LDISP_5975_ The SARS-CoV-2 S^B^ construct was synthesized by GenScript into pcDNA3.1-with an N-terminal mu-phosphatase signal peptide and a C-terminal hexahistidine tag (G-HHHHHH). The boundaries of the construct are N-terminal _337_RFPN_340_ and C-terminal _539_STNL_542_. Both constructs were produced in 500mL HEK293F cells grown in suspension using FreeStyle 293 expression medium (Life technologies) at 37°C in a humidified 8% CO_2_ incubator rotating at 130 rpm. The cultures were transfected using 293-Free transfection reagent (Millipore) with cells grown to a density of 1 million cells per mL and cultivated for 3 days. The supernatants were harvested and cells resuspended for another 3 days, yielding two harvests. Proteins were purified from clarified supernatants using a 5mL Cobalt affinity column (Takara), concentrated and flash frozen in a buffer containing 20 mM Tris pH 8.0 and 300 mM NaCl prior to analysis. SDS-PAGE was run to check purity.

### Transient expression of hACE2

Expression and purification of human angiotensin-converting enzyme ectodomain (ACE2, residues 1–614) fused to the Fc region of human IgG (hFc) was performed as previously described (Walls et al., 2019) and cleavage of the Fc fragment was carried out with thrombin. Briefly, hACE2 was produced in 500mL HEK293F cells grown in suspension using FreeStyle 293 expression medium (Life technologies) at 37°C in a humidified 8% CO_2_ incubator rotating at 130 rpm. The cultures were transfected using 293-Free transfection reagent (Millipore) with cells grown to a density of 1 million cells per mL and cultivated for 4 days. The supernatants were harvested and cells resuspended for another four days, yielding two harvests. Clarified supernatants were concentrated using Vivaflow tangential filtration cassettes (Sartorius, 10-kDa cut-off) before affinity purification using a Protein A column (GE LifeSciences) followed by gel filtration chromatography using a Superdex 200 10/300 GL column (GE Life Sciences) equilibrated in 20 mM Tris-HCl, pH 8, 100 mM NaCl. The Fc tag was removed by thrombin cleavage in a reaction mixture containing 3 mg of recombinant ACE2-FC ectodomain and 10 μg of thrombin in 10 mMTris-HCl pH8.0 and 20mM CaCl_2_.The reaction mixture was incubated at 25 °C overnight and re-loaded in a Protein A column to remove uncleaved protein and the Fc tag. The cleaved protein was further purified by gel filtration using a Superdex 200 column 10/300 GL (GE Life Sciences) equilibrated in a buffer containing 20 mM Tris pH 8.0 and 100 mM NaCl. The purified protein was quantified using absorption at 280 nm and concentrated to approximately 1 mg/ml.

### Pseudovirus production

MLV-based SARS-CoV S, SARS-CoV-2 S, and SARS-CoV-2 S_fur/mut_ pseudotypes were prepared as previously described (Millet and Whittaker, 2016). HEK293T cells were cotransfected using Lipofectamine 2000 (Life Technologies) with an S encoding-plasmid, an MLV Gag-Pol packaging construct and the MLV transfer vector encoding a luciferase reporter, according to the manufacturer’s instructions. Cells were incubated for 5 hours at 37°C with transfection medium. Cells were then washed with DMEM 2X and DMEM containing 10% FBS was added for 60 hours. The supernatants were then harvested and filtered through 0.45-mm membranes and concentrated with a 30kDa membrane for 10 minutes at 3,000 rpm and then frozen at −80°C.

### Western Blotting

SARS-CoV S, SARS-CoV-2 S, and SARS-CoV-2 S_Fur/Mut_ pseudovirions were thawed and 4X SDS loading buffer was added prior to boiling for 10 min at 95°C. Samples were run on a 4%–20% gradient Tris-Glycine Gel (BioRad) and transferred to a PVDF membrane. An anti-S_2_ SARS-CoV S monoclonal primary antibody (1:150 dilution) or an MLV p30 monoclonal antibody (1:1,000 dilution, AbCam) and an Alexa Fluor 680-conjugated goat anti-human or mouse secondary antibody (1:30,000 and 1:15,000 dilution respectively, Jackson Laboratory) were used for Western-blotting. A LI-COR processor was used to develop images.

### Pseudovirus entry assays

VeroE6 and Baby Hamster Kidney (BHK) cells were cultured in 10% FBS, 1% PenStrep DMEM. BHK or VeroE6 cells were plated into 12 well plates at a density of 0.3 x 10^6^ for 16 hours. BHK cells were either not transfected or transfected with 0.8 μg ACE2 per well using standard lipofectamine2000 (Life Technologies) protocols and incubated for another 16 hours. Concentrated pseudovirus was added to the wells after washing 3X with DMEM. After 2-3 hours 20% FBS and 2% PenStrep containing DMEM was added to the cells for 48 hours. Following the 48 hr infection, One-Glo-EX (Promega) was added to the cells in equivalent culturing volume and incubated in the dark for 10 minutes prior to reading on a Varioskan LUX plate reader (ThermoFisher). Measurements were done in triplicate and relative luciferase units (RLU) were plotted.

### Biolayer interferometry

Assays were performed on an Octet Red (ForteBio) instrument at 30°C with shaking at 1,000 RPM. Anti-penta His biosensors were hydrated in water for 30 minutes prior to a 1 min incubation in 10X kinetics buffer (undiluted). Either SARS-CoV-2 S^B^ or SARS-CoV S^B^ were loaded at 10-20 μg/mL in 10X Kinetics Buffer for 200-900 seconds prior to baseline equilibration for 180 seconds in 10X kinetics buffer. Association in 10X kinetics buffer diluted ACE2 at various concentrations in a two-fold dilution series from 20nM to 2.5nM was carried out for 1,000 seconds prior to dissociation at 10X kinetics buffer for 1,000 seconds. The data were baseline subtracted prior to fitting performed using a 1:1 binding model and the ForteBio data analysis software. Mean k_on_, k_off_ values were determined with a global fit applied to all data. The experiments were done with two separate purification batches of SARS-CoV S^B^, SARS-CoV-2-S^B^, and ACE2 in triplicate and the average is reported.

### Protein expression and purification

The SARS-CoV-2 2P S and SARS-CoV 2P S ectodomains were produced in 500mL HEK293F cells grown in suspension using FreeStyle 293 expression medium (Life technologies) at 37°C in a humidified 8% CO2 incubator rotating at 130 r.p.m. The cultures were transfected using 293fectin (ThermoFisher Scientific) with cells grown to a density of 1 million cells per mL and cultivated for three days. The supernatants were harvested and cells resuspended for another three days, yielding two harvests. Clarified supernatants were purified using a 5mL Cobalt affinity column (Takara). Purified protein was filtered or concentrated and flash frozen in Tris-saline (50 mM Tris pH 8.0, 150 mM NaCl) prior to negative staining and cryo-EM analysis.

### Cryo-EM sample preparation and data collection

3 μL of 0.16 mg/mL SARS-CoV-2 S was loaded onto a freshly glow discharged (30 s at 20 mA) lacey carbon grid with a thin layer of evaporated continuous carbon prior to plunge freezing using a vitrobot MarkIV (ThermoFisher Scientific) using a blot force of −1 and 2.5 second blot time at 100% humidity and 25°C. Data were acquired using the Leginon software (Suloway et al., 2005) to control an FEI Titan Krios transmission electron microscope operated at 300 kV and equipped with a Gatan K2 Summit direct detector and Gatan Quantum GIF energy filter, operated in zero-loss mode with a slit width of 20 eV. Automated data collection was carried out using Leginon at a nominal magnification of 130,000x with a pixel size of 0.525 Å. The dose rate was adjusted to 8 counts/pixel/s, and each movie was acquired in super-resolution mode fractionated in 50 frames of 200 ms. 3,000 micrographs were collected in a single session with a defocus range comprised between 1.0 and 2.5 μm.

### CryoEM data processing

Movie frame alignment, estimation of the microscope contrast-transfer function parameters, particle picking and extraction were carried out using Warp (Tegunov and Cramer, 2019). Particle images were extracted with a box size of 800 binned to 400 yielding a pixel size of 1.05 Å. For each data set two rounds of reference-free 2D classification were performed using cryoSPARC (Punjani et al., 2017) to select well-defined particle images. Subsequently, two rounds of 3D classification with 50 iterations each (angular sampling 7.5°for 25 iterations and 1.8° with local search for 25 iterations) were carried out using Relion (Zivanov et al., 2018) without imposing symmetry to separate distinct SARS-CoV-2 S conformations. 3D refinements were carried out using non-uniform refinement in cryoSPARC (Punjani et al., 2017) using an ab initio model generated using cryoSPARC. Particle images were subjected to Bayesian polishing (Zivanov et al., 2019) before performing another round of on-uniform refinement in cryoSPARC (Punjani et al., 2017) followed by per-particle defocus refinement and again non-uniform refinement. Local resolution estimation, filtering and sharpening was carried out using CryoSPARC. Reported resolutions are based on the gold-standard Fourier shell correlation (FSC) of 0.143 criterion and Fourier shell correlation curves were corrected for the effects of soft masking by high-resolution noise substitution (Scheres and Chen, 2012).

### CryoEM model building and analysis

UCSF Chimera (Goddard et al., 2007) and Coot were used to fit atomic models (PDB 6NB6) into the cryoEM map. The model was subsequently manually rebuilt using Coot and the C3-symmetrized map (Brown et al., 2015; Emsley et al., 2010). This model was then used to build features specific of the C1-map. N-linked glycans were hand-built into the density where visible and the models were refined and relaxed using Rosetta (Wang et al., 2016). Glycan refinement relied on a dedicated Rosetta protocol, which uses physically realistic geometries based on prior knowledge of saccharide chemical properties (Frenz, 2019), and was aided by using both sharpened and unsharpened maps. Models were analyzed using MolProbity (Chen et al., 2010) and privateer (Agirre et al., 2015). Figures were generated using UCSF ChimeraX (Goddard et al., 2018).

### Immunizations with SARS-CoV S

57BL/6J mice were obtained from the Jackson Laboratory and maintained at the Comparative Medicine Facility at the Fred Hutchinson Cancer Research Center. After collecting a pre-bleed, mice were immunized with 5μg of SARS-CoV 2P S ectodomain trimer formulated in 50μl of PBS and 50μl of Sigma Adjuvant System subcutaneously in the base of the tail at weeks 0, 4, and 12 and bleeds were collected 2 weeks after each immunization. Plasma from immunized animals was heat inactivated at 56°C for 1 hour and then stored at 4°C until use. All experiments were conducted at the Fred Hutchinson Cancer Research Center according to approved Institutional Animal Care and Use Committee protocols.

