## Supplemental material for "Structure, function and antigenicity of the SARS-CoV-2 spike glycoprotein"

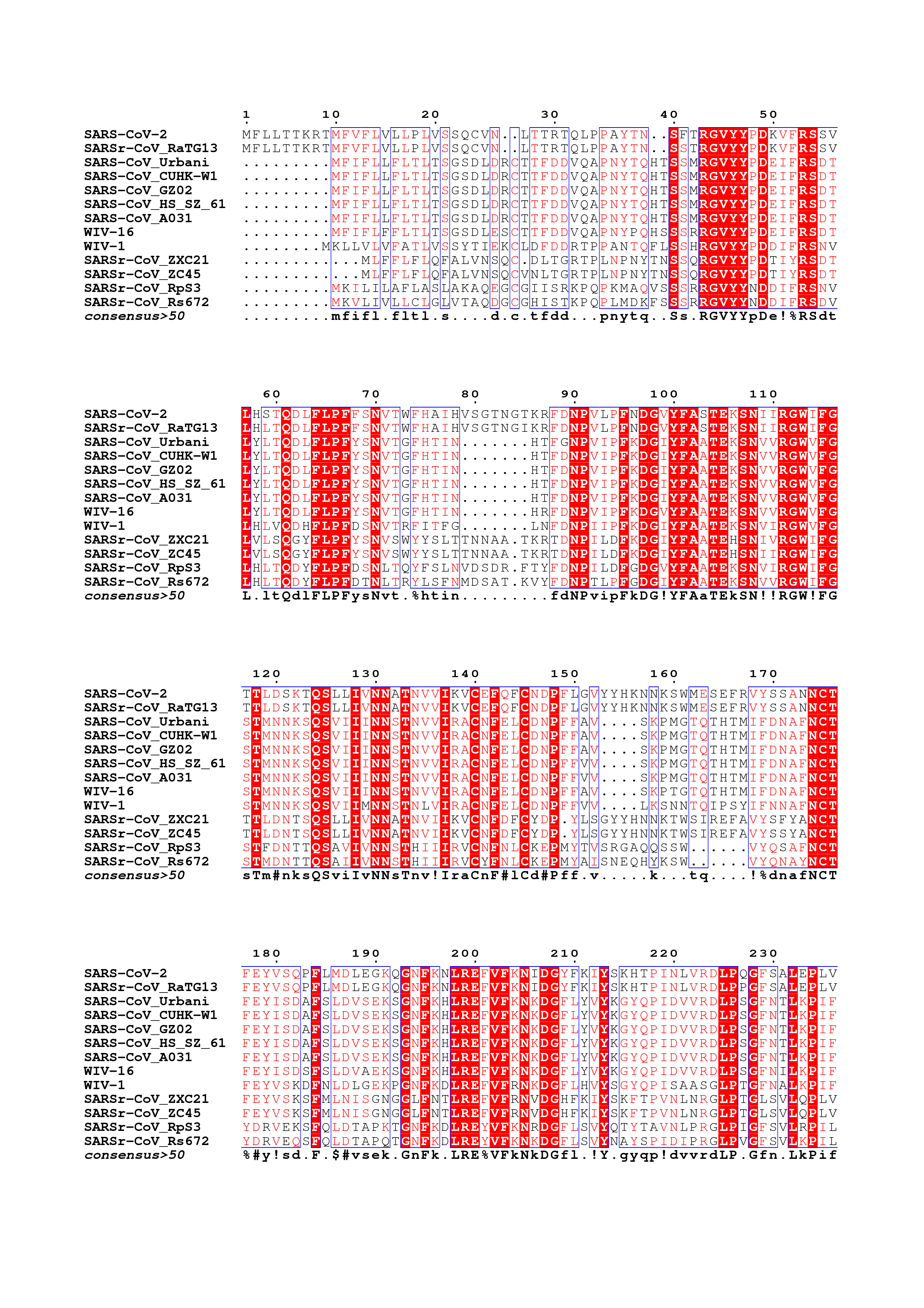

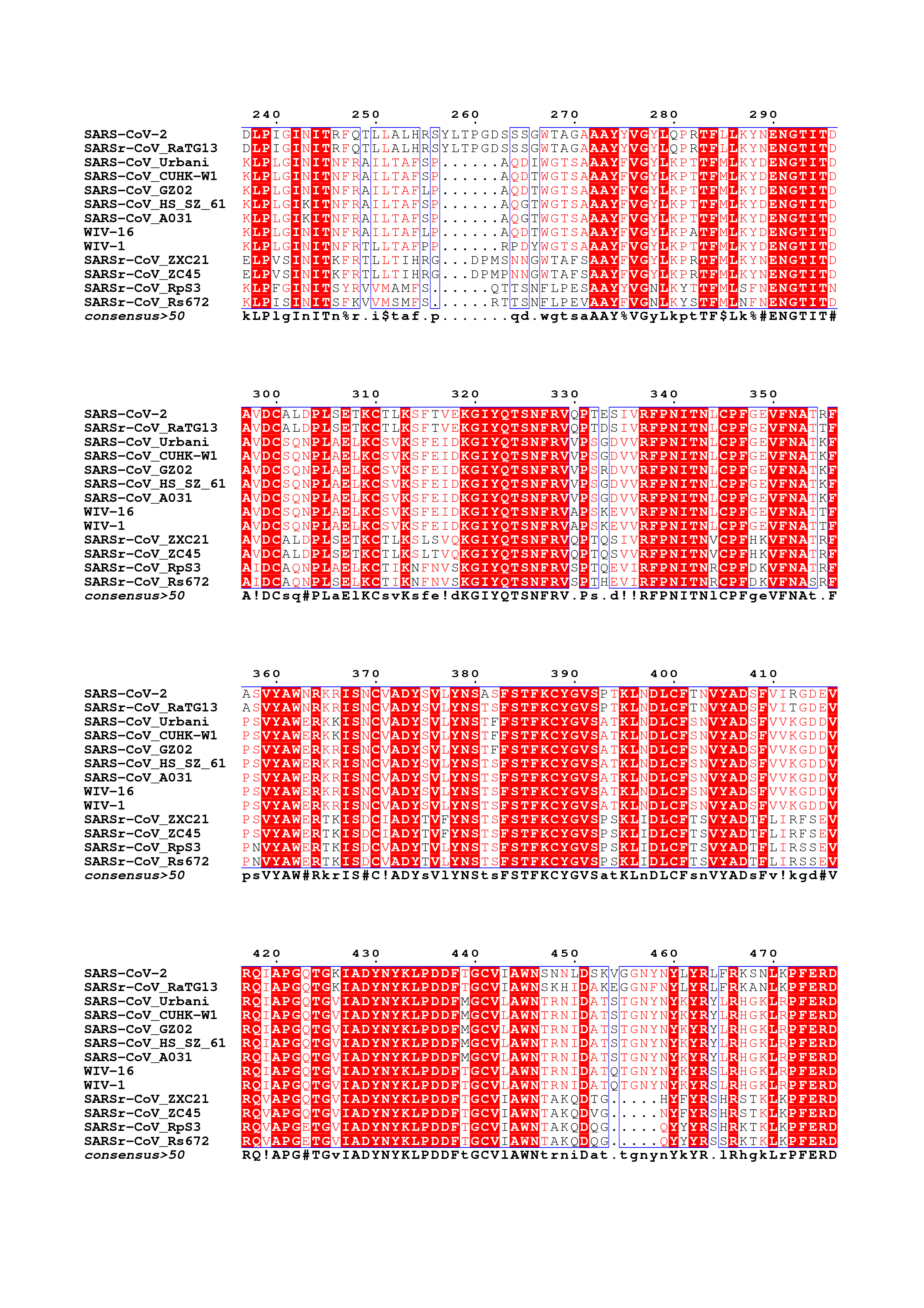

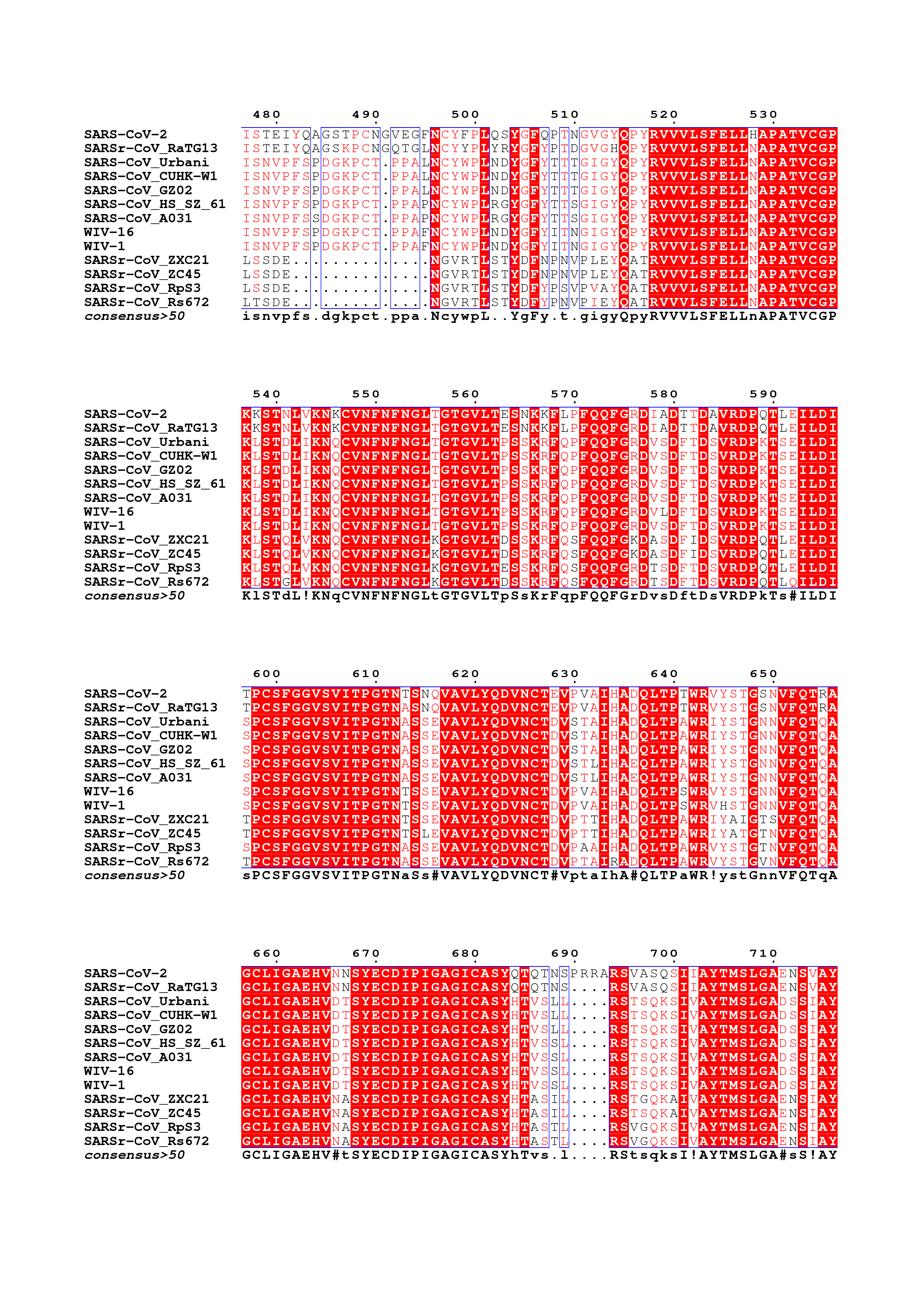

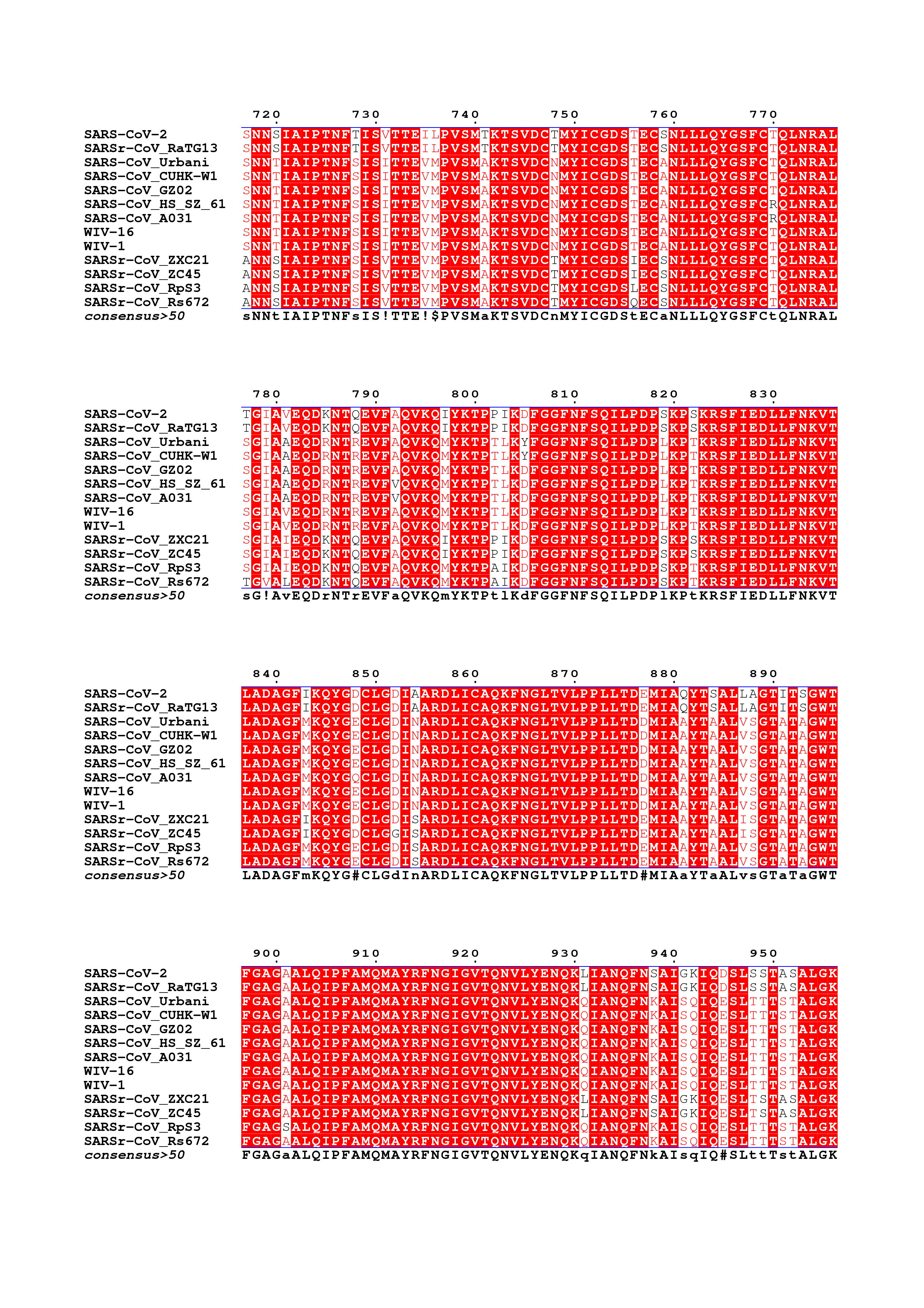

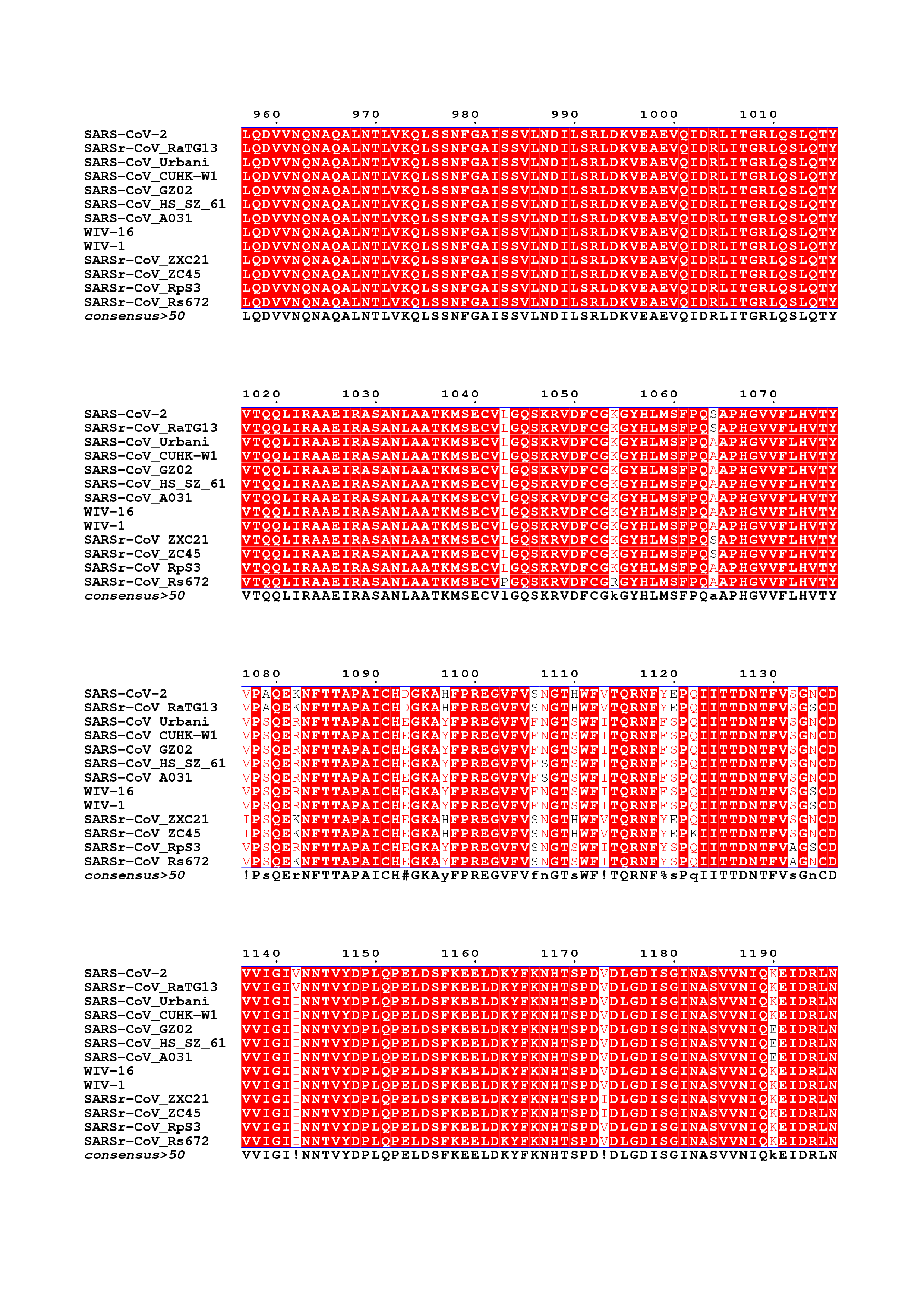

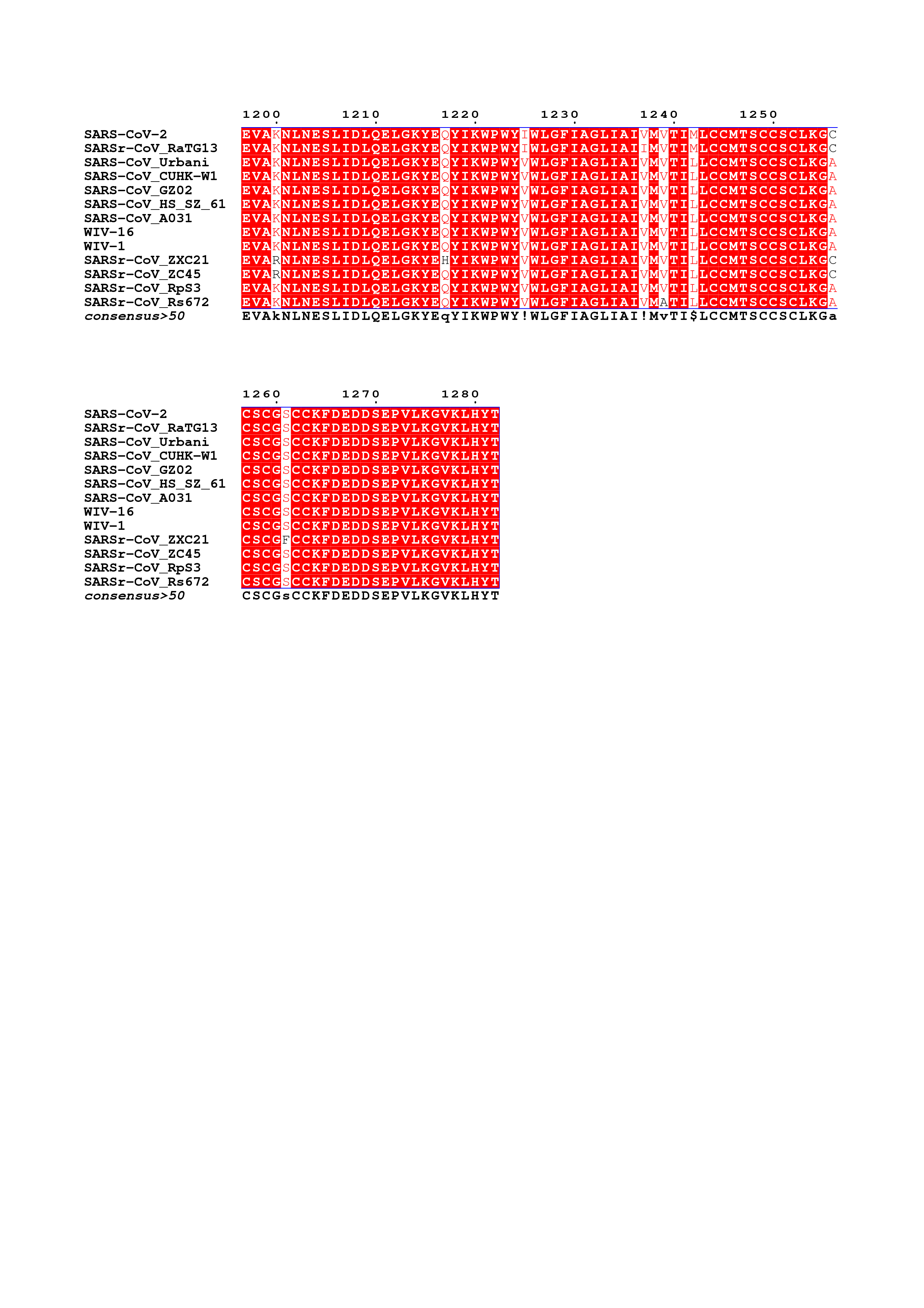

**Figure S1. Amino acid sequence alignment of sarbecovirus S glycoproteins.** The following sequences were used: SARS-CoV-2 (), SARSr-CoV RaTG13 (QHR63300.1), SARS-CoV Urbani (AAP13441.1), SARS-CoV CUHK-W1 (AAP13567.1), SARS-CoV GZ02 (AAS00003.1), SARS-CoV HS/SZ/61/03 (AY515512.1), SARS-CoV A031 (AAV97988.1), WIV-16 (ALK02457.1), WIV-1 (AGZ48828.1), SARSr-CoV ZXC21 (AVP78042.1), SARSr-CoV ZC45 (AVP78031.1), SARSr-CoV Rp3 (Q3I5J5.1), SARSr-CoV Rs672 (ACU31032.1).

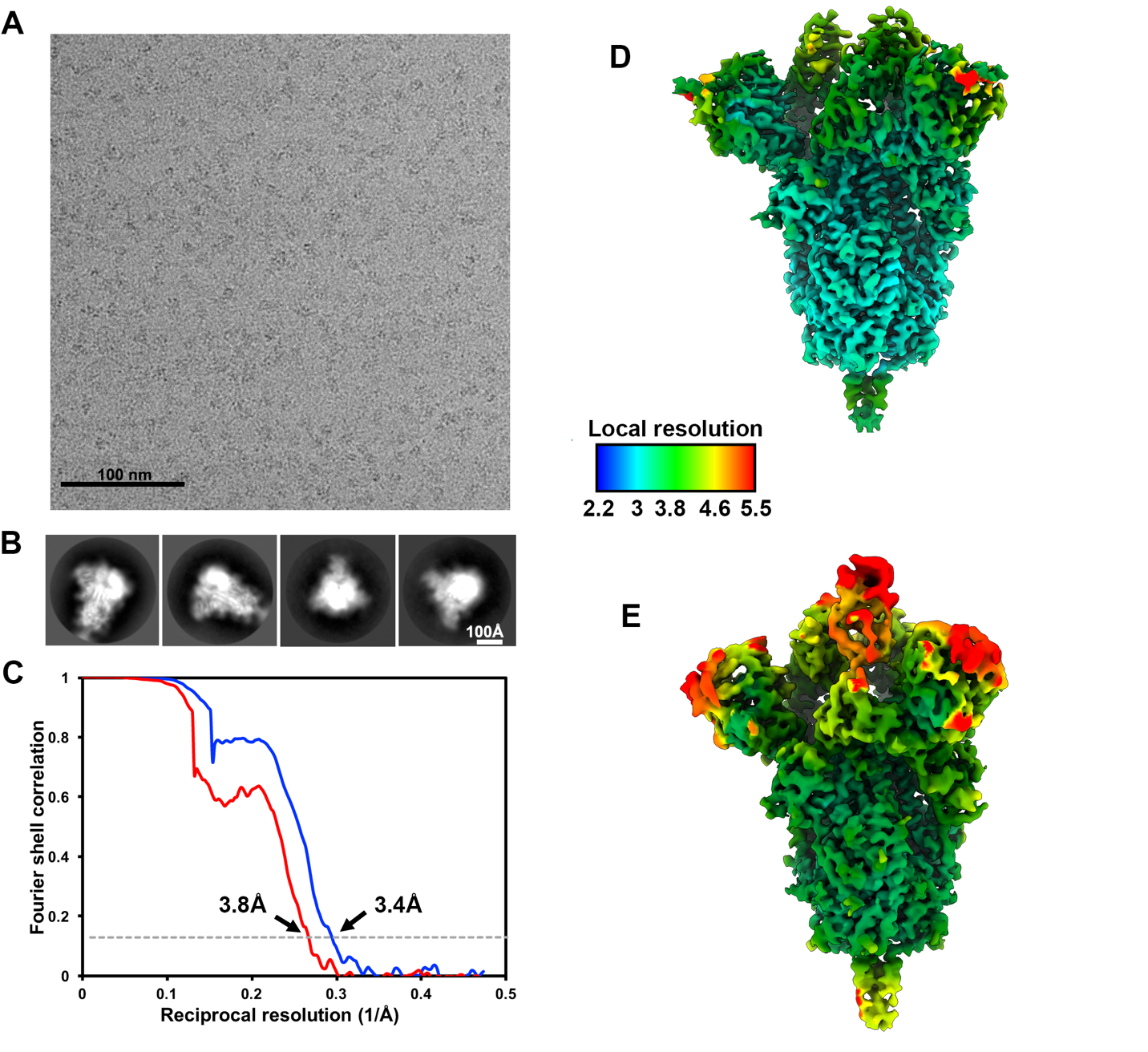

**Figure S2. CryoEM data processing and validation. A-B.** Representative electron micrograph (A) and class averages (B) of SARS-CoV-2 S embedded in vitreous ice. **C.** Gold-standard Fourier shell correlation curves for the closed (blue) and partially open trimers (red). The 0.143 cutoff is indicated by horizontal dashed lines. **D-E.** Local resolution map calculated using cryoSPARC for the closed (D) and partially open (E) reconstructions.

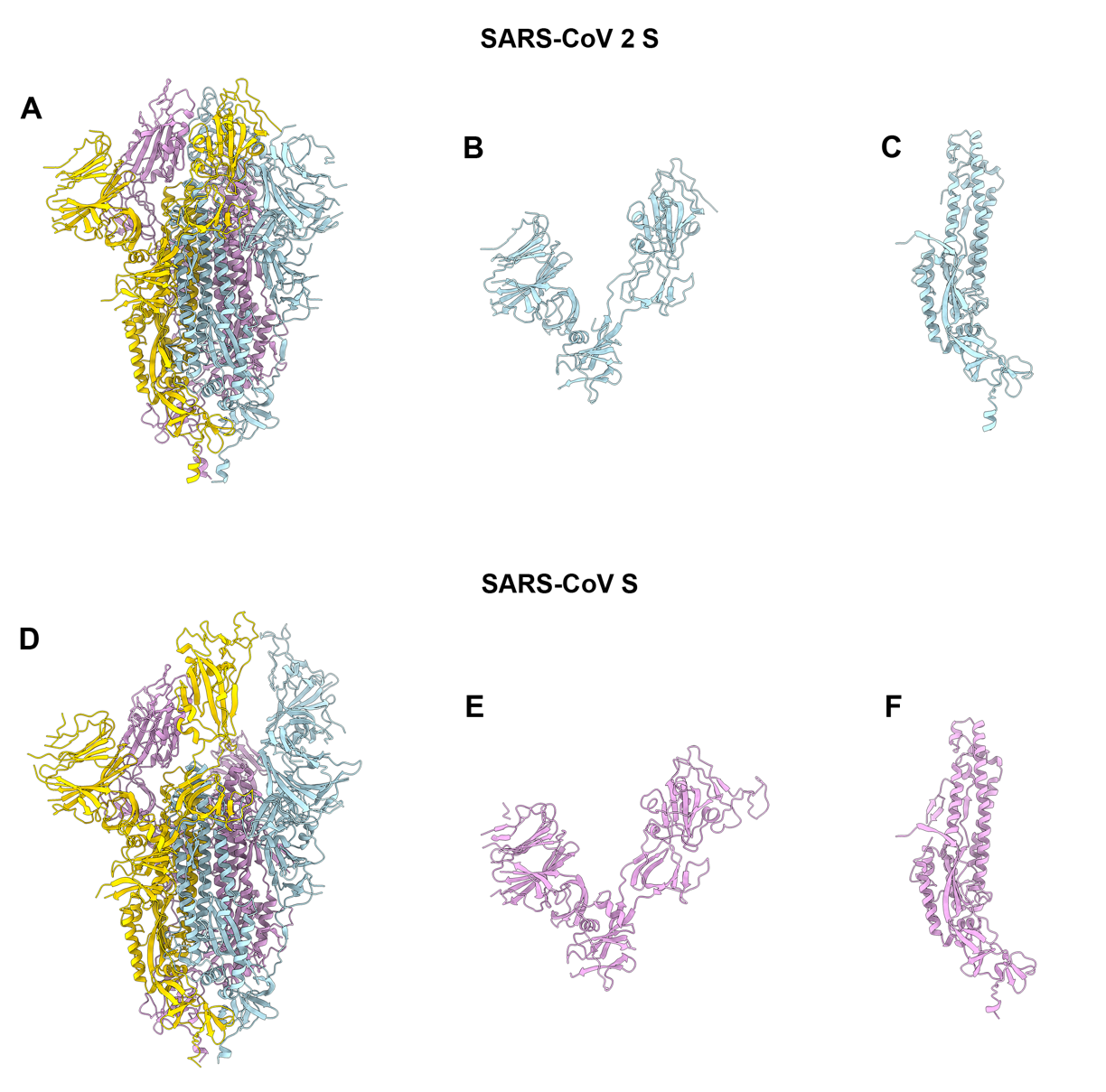

**Figure S3. Comparison of the SARS-CoV-2 and SARS-CoV S structures. A-D.** Ribbon diagrams of the SARS-CoV-2 S (A) and SARS-CoV S (PDB 6NB6, D) ectodomain cryoEM structures. **B-E,** The SARS-CoV-2 (C) and SARS-CoV (E) S_1_ subunits. **C-F,** The SARS-CoV-2 (C) and SARS-CoV (F) S_2_ subunits.

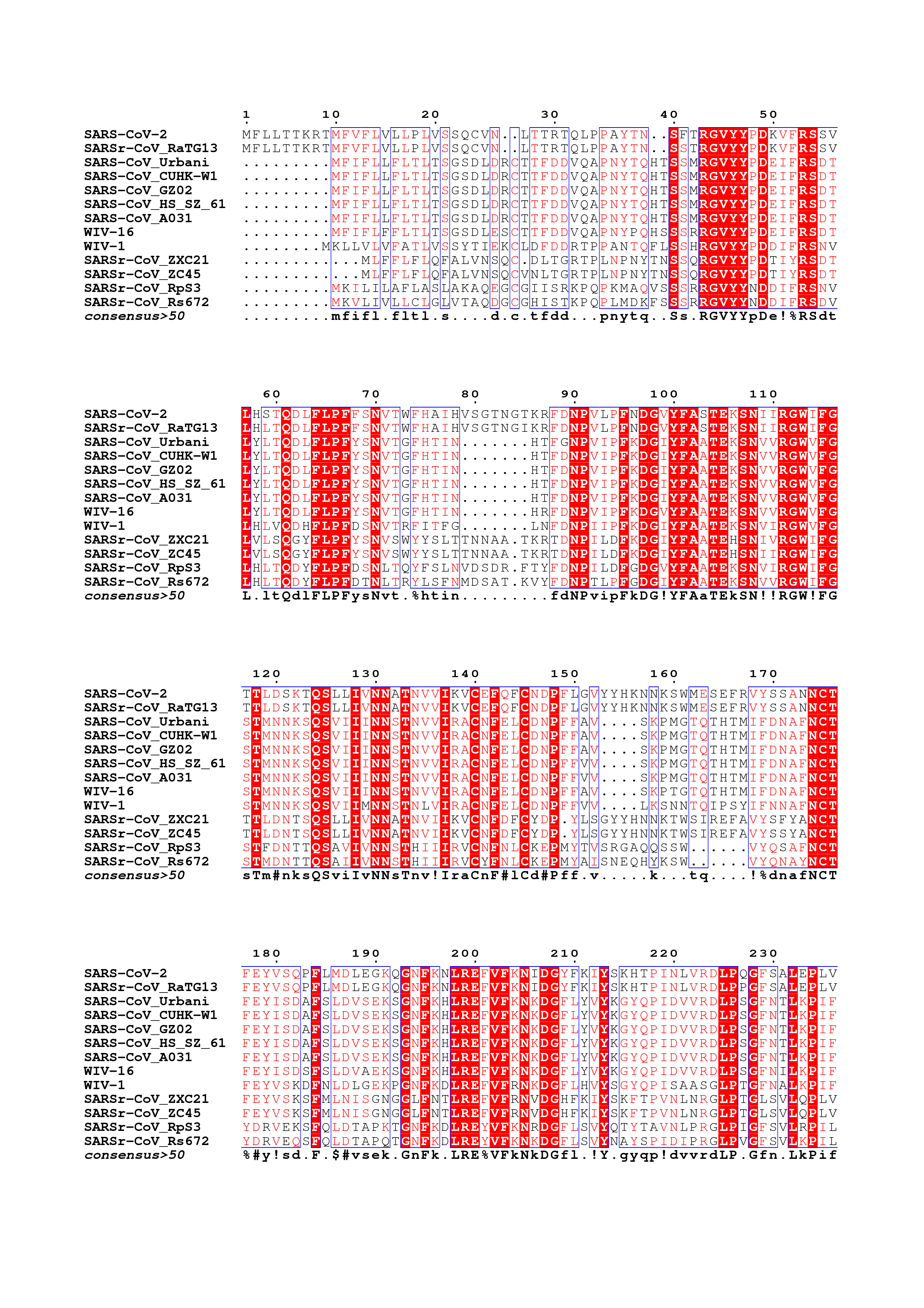

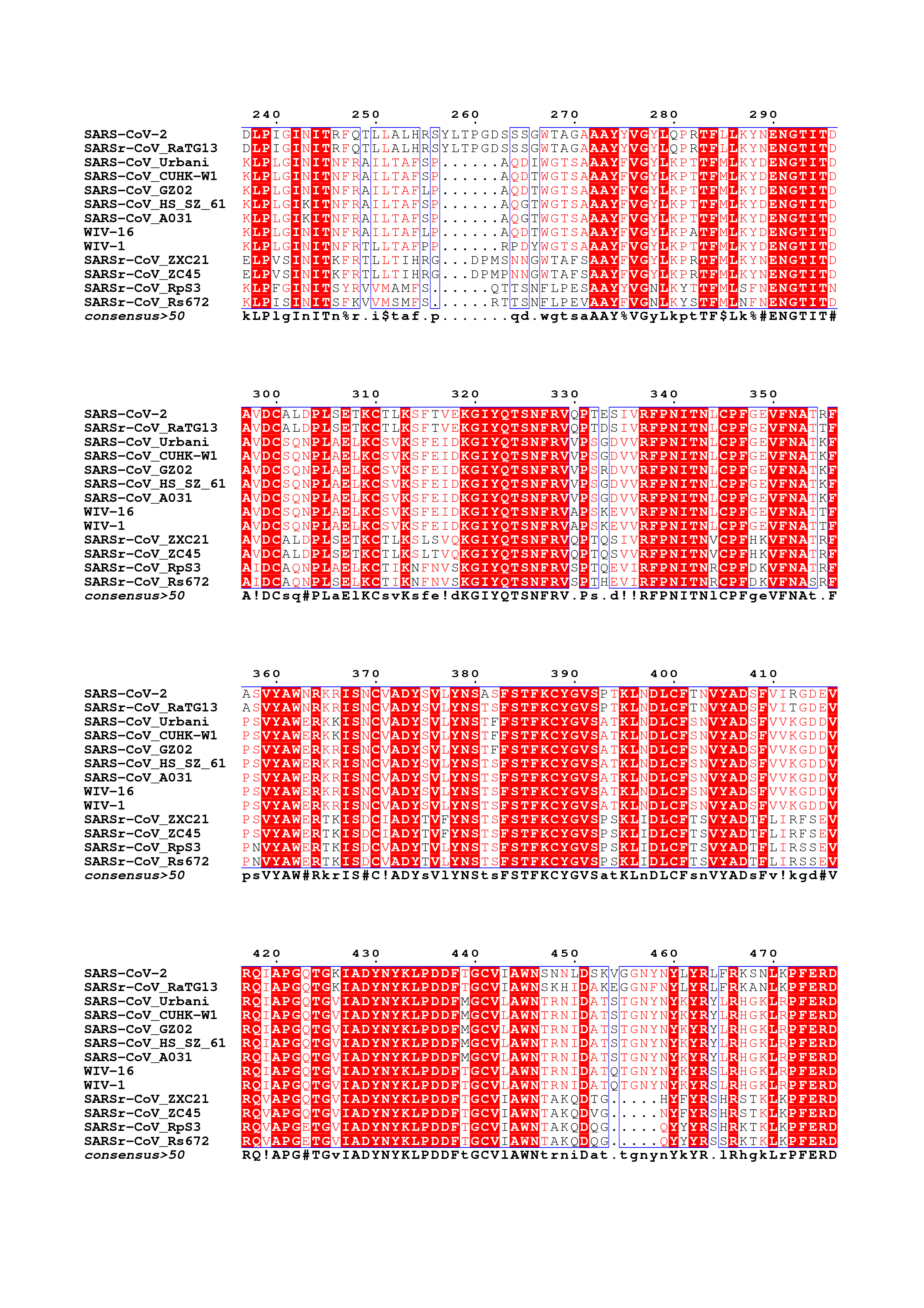

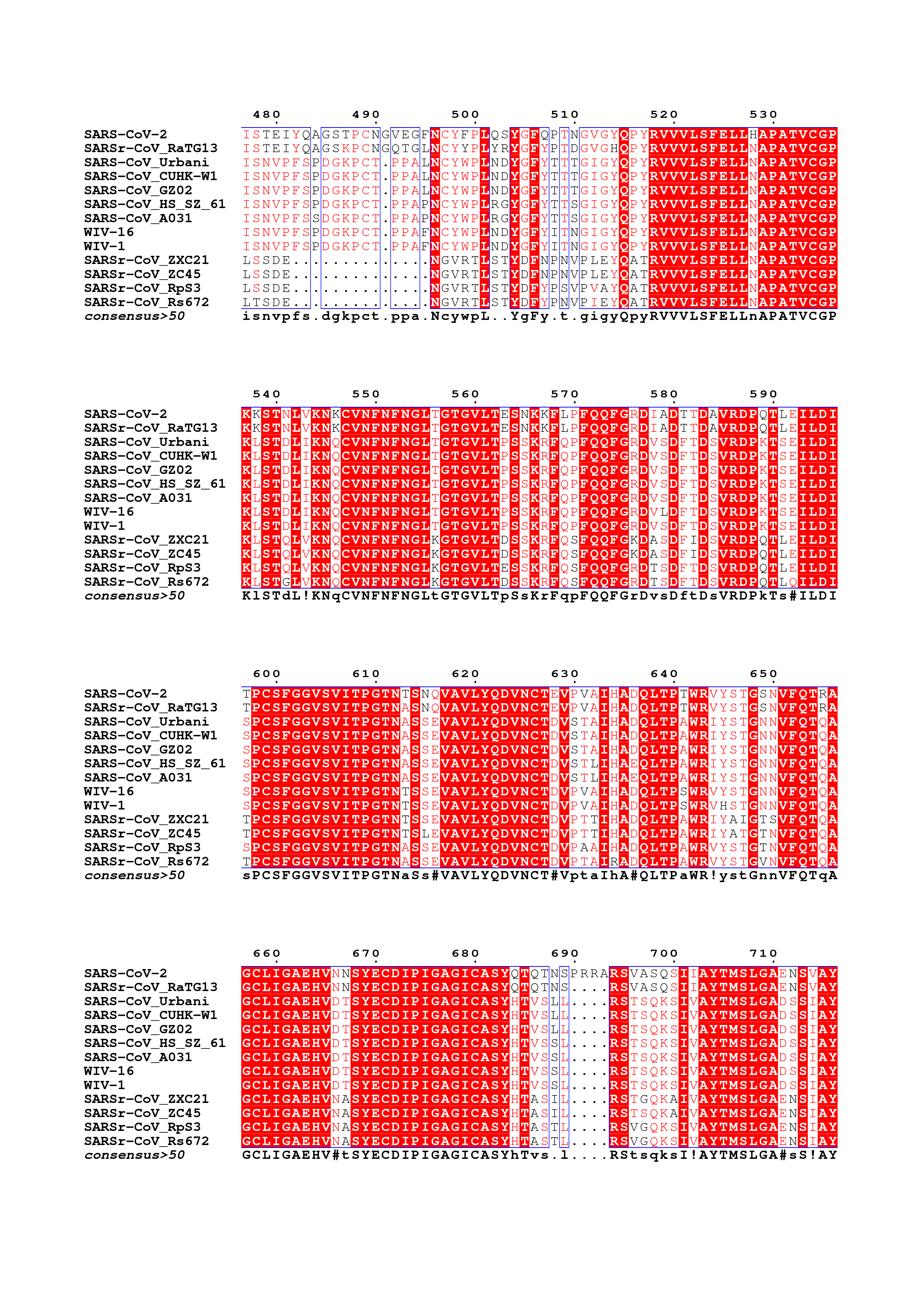

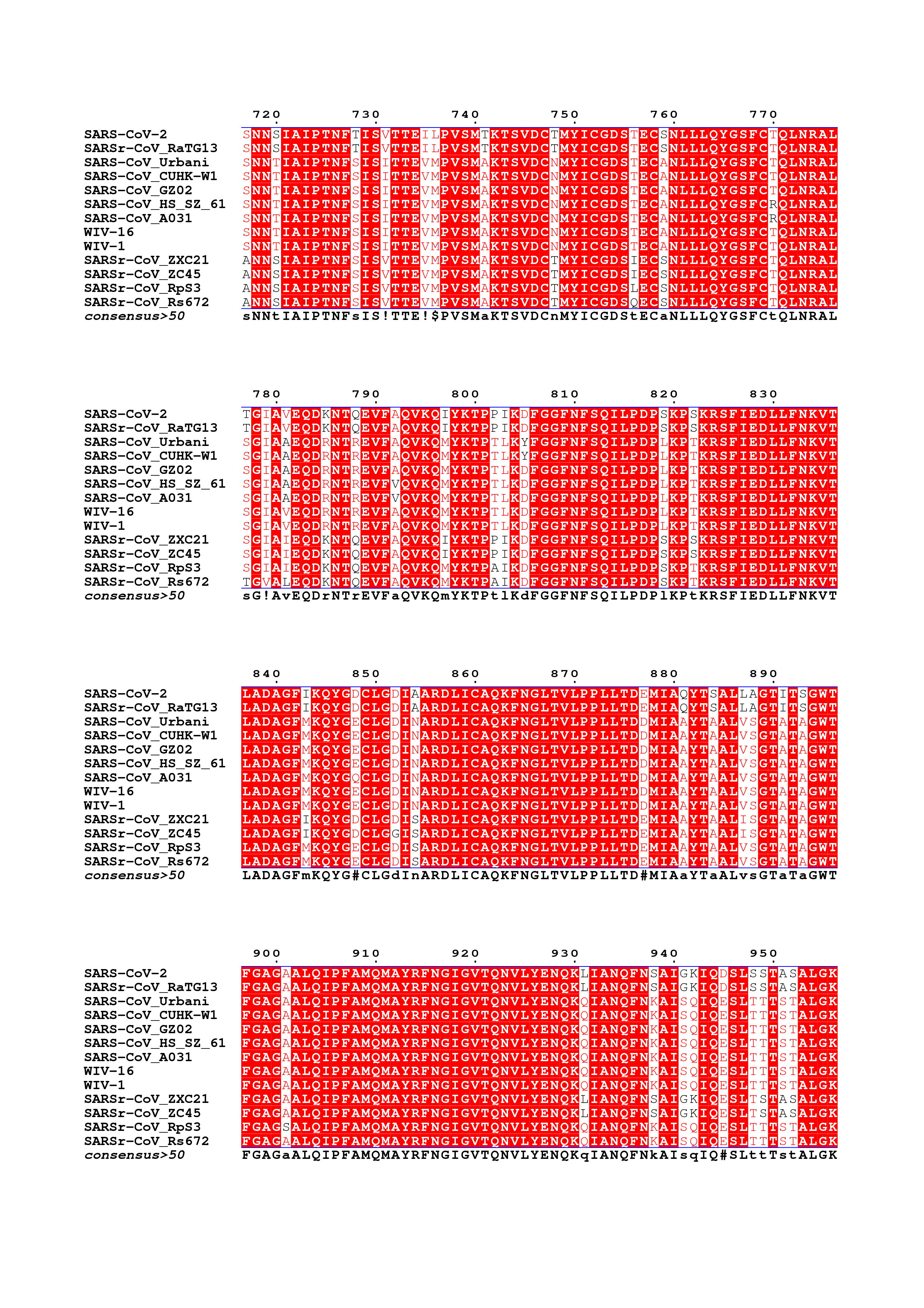

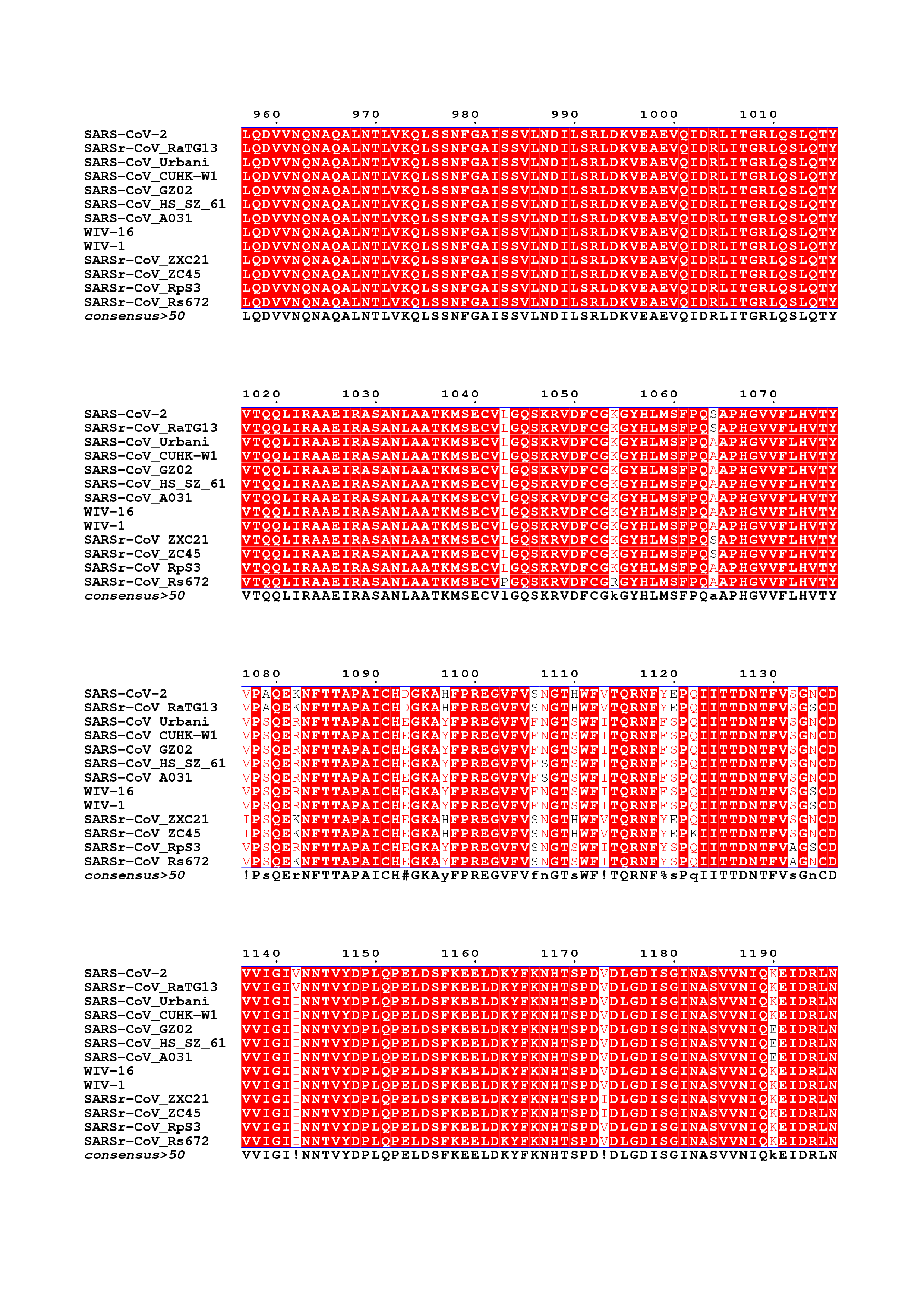

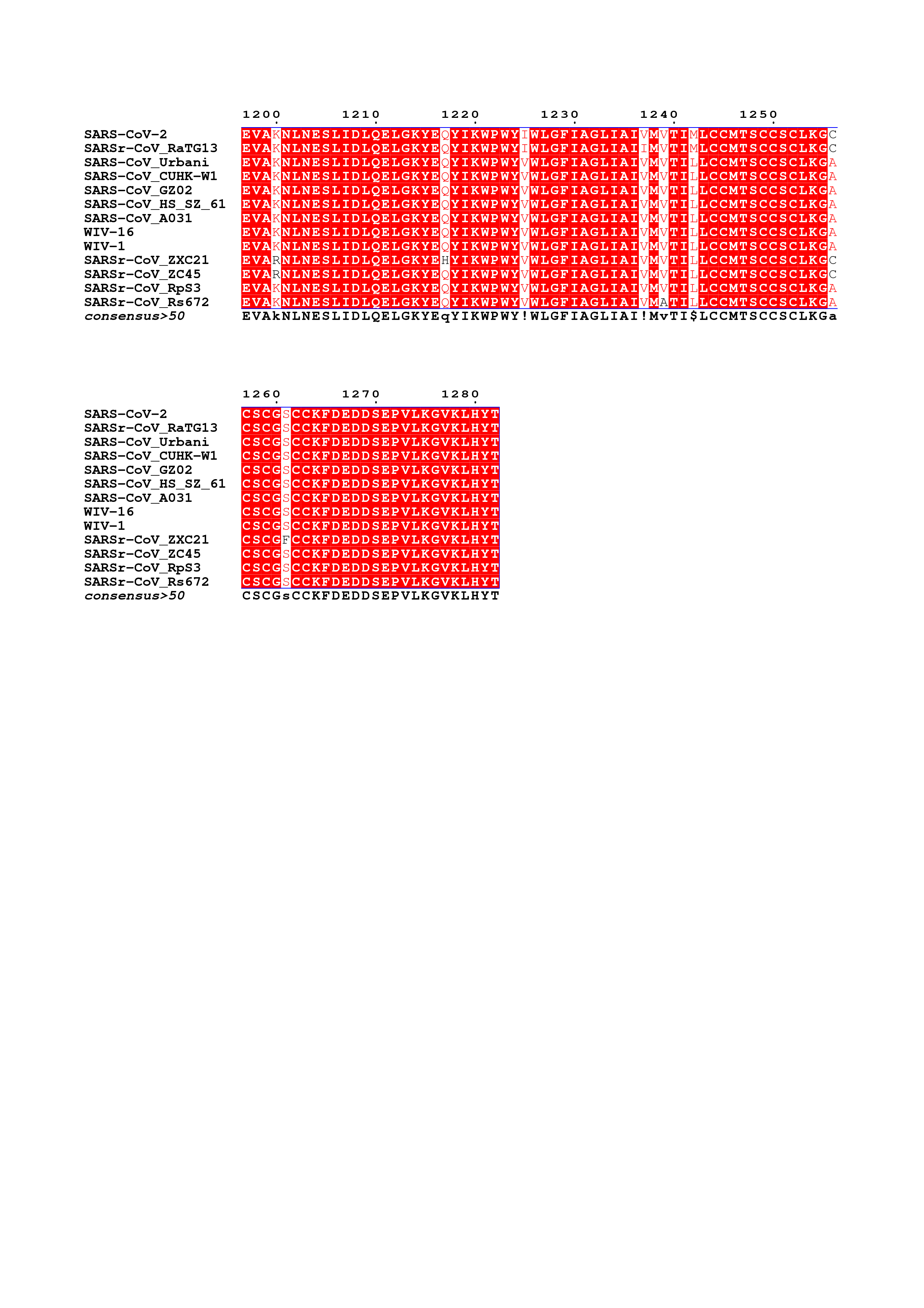

**Figure S1. Amino acid sequence alignment of sarbecovirus S glycoproteins.** The following sequences were used: SARS-CoV-2 (), SARSr-CoV RaTG13 (QHR63300.1), SARS-CoV Urbani (AAP13441.1), SARS-CoV CUHK-W1 (AAP13567.1), SARS-CoV GZ02 (AAS00003.1), SARS-CoV HS/SZ/61/03 (AY515512.1), SARS-CoV A031 (AAV97988.1), WIV-16 (ALK02457.1), WIV-1 (AGZ48828.1), SARSr-CoV ZXC21 (AVP78042.1), SARSr-CoV ZC45 (AVP78031.1), SARSr-CoV Rp3 (Q3I5J5.1), SARSr-CoV Rs672 (ACU31032.1).

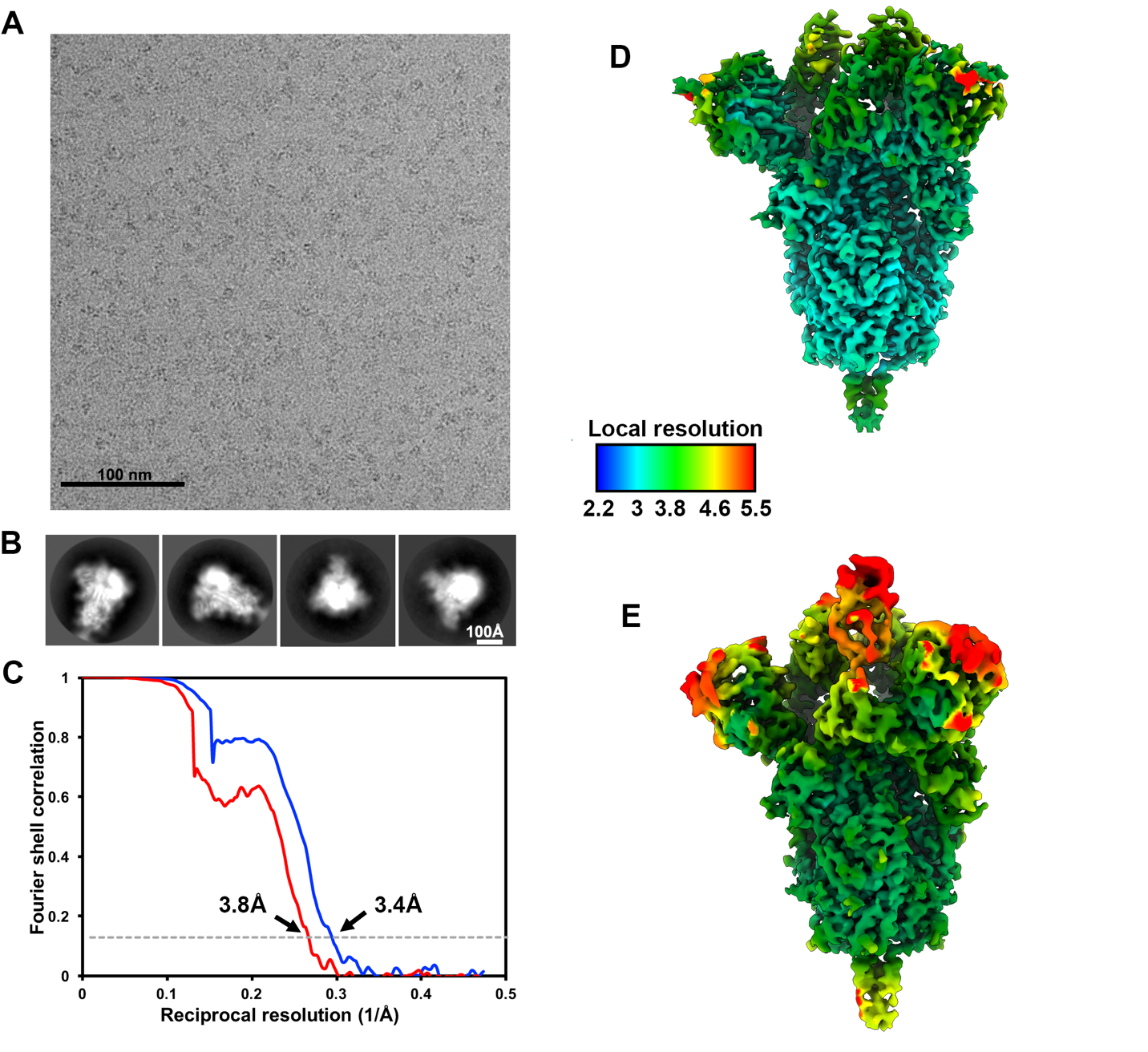

**Figure S2. CryoEM data processing and validation. A-B.** Representative electron micrograph (A) and class averages (B) of SARS-CoV-2 S embedded in vitreous ice. **C.** Gold-standard Fourier shell correlation curves for the closed (blue) and partially open trimers (red). The 0.143 cutoff is indicated by horizontal dashed lines. **D-E.** Local resolution map calculated using cryoSPARC for the closed (D) and partially open (E) reconstructions.

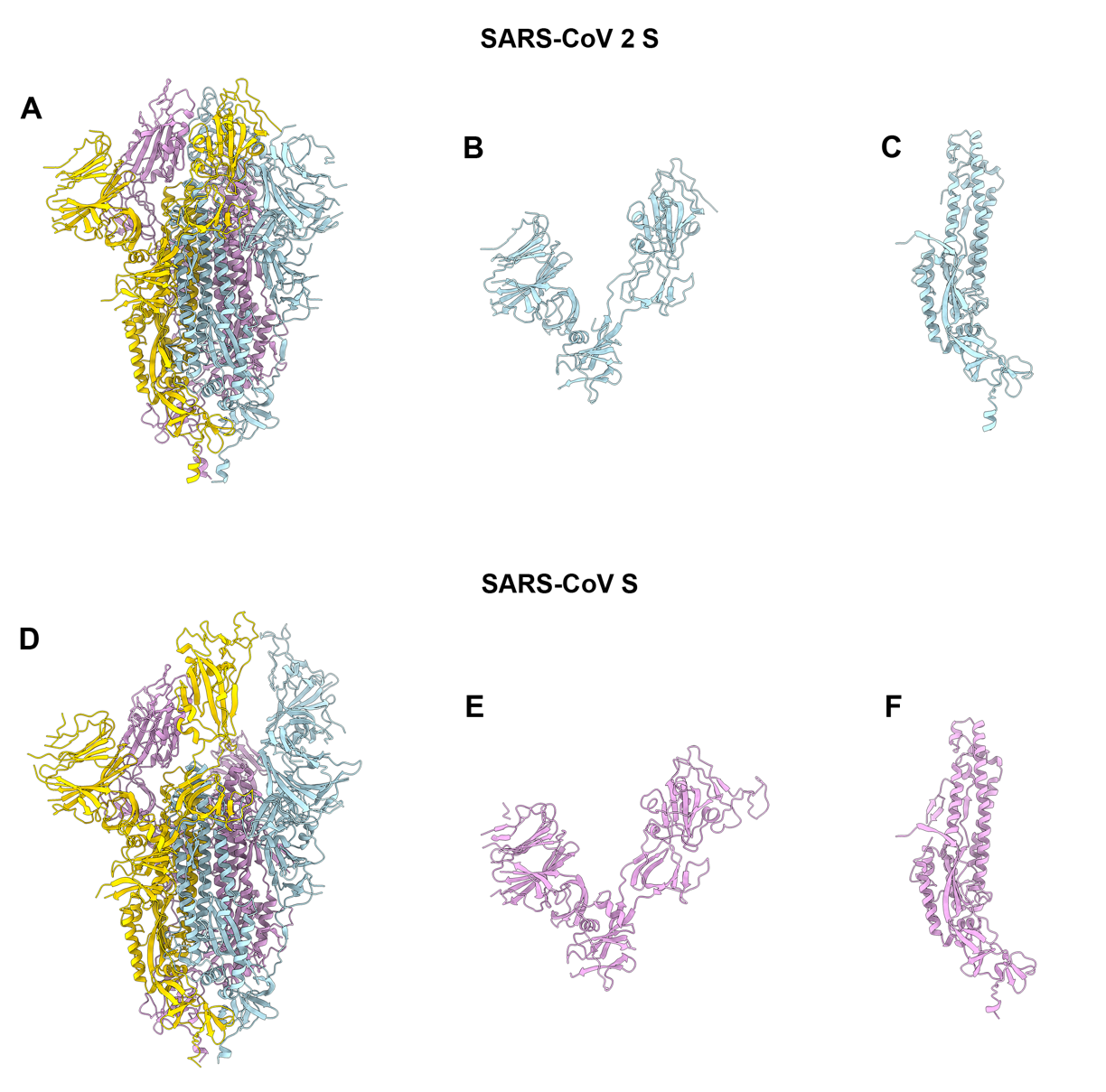

**Figure S3. Comparison of the SARS-CoV-2 and SARS-CoV S structures. A-D.** Ribbon diagrams of the SARS-CoV-2 S (A) and SARS-CoV S (PDB 6NB6, D) ectodomain cryoEM structures. **B-E,** The SARS-CoV-2 (C) and SARS-CoV (E) S_1_ subunits. **C-F,** The SARS-CoV-2 (C) and SARS-CoV (F) S_2_ subunits.
